# Joint Walking Training with Virtual and Real Partners: Testing the Transfer from Virtual Reality to the Real World

**DOI:** 10.64898/2026.11.25.767275

**Authors:** Constanze Dammeyer, Mathilde Truffer, Damian Beck, Ralf Kredel, Stephan Zahno

## Abstract

**Background:** Joint walking, in which two people continuously coordinate their steps, has been proposed as a promising approach to improve gait in older adults. Virtual reality (VR) offers new possibilities for joint walking training, including the potential to practise with a virtual partner. However, it remains unclear whether learning to coordinate with a partner in VR transfers to the real world. Here, as a first step towards VR-based training in older adults and clinical populations, we investigate learning and transfer effects of joint walking training in real and virtual environments in healthy young adults.

**Methods:** In a pre-post design, 36 participants practised following a leader’s rhythmic forward and backward steps (1 Hz) to music while keeping hand-to-chest contact. Participants trained either with a real partner, with the partner visually presented as an avatar through a head-mounted display or walked to the beat without a partner. Coordination performance was quantified as the dispersion of the temporal difference between the leader’s and follower’s chest acceleration maxima and assessed before and after training in both the real and virtual environment. Linear mixed models were used to test the effects of time (pre, post) and training group (virtual, real, solo) in both test environments (real, virtual).

**Results:** From pre- to post-test, all training groups showed improved leader–follower coordination, reflected by reduced dispersion of the temporal difference, in both the virtual (*b* = −14.46, 95% CI [−25.85, −3.06], *t*(35) = −2.58, *p* = .014) and real test environments (*b* = −26.78, 95% CI [−37.09, −16.47], *t*(35) = −5.27, *p* < .001). These improvements did not differ between training groups.

**Conclusions:** Our findings indicate that VR training transfers to the real world, supporting the potential of VR joint walking training as a promising approach for gait training in older adults. However, since improvements were also observed after solo walking, specific transfer effects from joint walking in VR to the real world should be investigated in future studies, particularly under more challenging test conditions and with older adults.

## Background

Ageing is associated with declines in gait and mobility [1], which increase the risk of falls and compromise a person’s independence and quality of life [2]. A particular challenge in everyday life is maintaining stable and safe gait while simultaneously adapting to unexpected changes in the environment [3].

One promising approach for gait training addressing continuous adaptations is walking together with another person. In a systematic review, Felsberg and Rhea [4] found that side-by-side walking and walking with hand contact can induce interpersonal gait synchronization and highlighted the potential of joint walking for gait rehabilitation. As possible mechanisms for these gait changes, the authors suggest sensory feedback (especially touch and vision), and the stabilizing influence of a healthy partner’s gait pattern [4]. Individual studies indicate that interaction with a walking partner can modify specific aspects of gait [5–8]. For example, walking with a stable partner improved gait symmetry and bilateral coordination [9, 10]. Beyond immediate adaptations, Almurad et al. [11] showed that prolonged arm-in-arm walking with a younger partner restores long-range correlations in stride fluctuations among older adults, a pattern associated with healthier gait dynamics and lower fall risk. A recent study [12] showed that three weeks of arm-in-arm walking training with a younger partner reduced fear of falling in older adults and altered the temporal structure of gait fluctuations, with the latter effect persisting for up to two months after training. Together, these findings suggest that joint walking may provide a promising approach for gait training in older adults.

Recent advances in virtual reality (VR) technology have created novel opportunities for research [13–16] and practice in rehabilitation settings [17–20]. From a research perspective, a major advantage of VR is that it enables the study of complex and naturalistic behaviour in an experimentally controlled manner [17, 19, 21]. For practice, VR-based training and therapy can provide real-time feedback, expose individuals to consistent or systematically varied conditions, and allow intervention parameters, such as pacing and intensity, to be flexibly adapted to individual needs [18, 19]. Specifically, for joint walking training, VR might offer the possibility of practising gait coordination with a virtual partner without requiring the presence of another person. As learning to coordinate with others often depends on the availability of training partners, virtual partners may help overcome this practical limitation [22]. Such independence would be particularly valuable for rehabilitation, as training could be offered more flexibly and even at home [17].

However, the practical value of VR-based training critically depends on whether the acquired skills transfer to real-world performance [23–28]. Yet, empirical support for transfer of sensorimotor learning from virtual to real-world environments remains limited. For sports, a systematic review by Michalski et al. [29] identified only four studies systematically testing VR to real-world transfer with both a control group and a measure of real-world transfer. While the findings from these four articles are encouraging and provide initial support for the transfer of VR-based skills training to real-world performance, Richlan et al. [30] noted that the number of intervention studies remains low and that systematic evidence on the factors underlying real-world transfer is still lacking. Regarding interpersonal coordination in VR, Varlet et al. [22] found that participants who practised with a virtual rowing partner improved their synchronization with a real teammate afterwards, showing that VR-based training of interpersonal coordination can transfer to real-world dyadic interaction.

Focusing on gait training in VR, systematic reviews have shown that VR-based training can improve balance and gait in older adults [20, 31], highlighting the potential of VR training to improve gait performance beyond the virtual environment itself. Training with virtual obstacles on a treadmill improved obstacle crossing in real-world overground walking in healthy young adults [23] and led to fewer falls in older adults at high risk of falls than treadmill training alone [32]. However, these studies concern solo gait training rather than joint walking. Initial studies investigating coordination with virtual partners have shown that participants can adapt their stepping timing to perturbations of a virtual avatar [33] and synchronize their steps with an avatar at levels comparable to those observed with a real person [34]. To the best of our knowledge, only one study has investigated gait training with a virtual partner in older adults [35]. Using augmented reality, Miller et al. [35] showed that gradually manipulating the distance and timing of a virtual walking partner altered their stride length, cycle time, and gait speed. While these findings provide initial evidence for carry-over effects of virtual-partner-based gait guidance, effects of practising walking with a virtual partner on subsequent interpersonal coordination with a real person remain unexplored.

In the current study, we address this gap by investigating the effects of joint walking training in a real and a virtual environment and whether learning transfers between environments. We developed a VR version of a joint walking task that Truffer et al. [36] previously used to experimentally study interpersonal coordination. In this task, participants act as followers and stand face-to-face with a leader, connected by a hand-to-chest contact. The leader is visually perceived either in the real world or as an avatar presented through a head-mounted display (HMD) in a virtual environment. The task is to coordinate forward and backward steps with the leader. Critically, the sequence of the leader’s forward and backward steps is unknown to the participants, creating uncertainty and requiring them to continuously adapt their steps based on incoming multimodal sensory information. We tested young, healthy adults to assess the general learning and VR to real-world transfer effects, with the long-term goal of applying this task as a training tool for older adults or in rehabilitation contexts. The task may be particularly relevant for such applications as it combines interpersonal coordination with the continuous need to adapt stepping under uncertainty, thereby targeting adaptive gait control, a capacity relevant to everyday mobility [3].

In a pre-post design, participants practised the joint walking task with a partner either in a real or a virtual environment or, as a control condition, walked to a beat without a partner. Before and after training, we assessed interpersonal coordination performance in the real and the virtual environment. We expected that (1) both joint-walking training groups would show greater pre- to-post improvements in coordination performance than the control group and (2) participants trained in the virtual environment would also improve their coordination in the real environment, indicating VR-to-real-world transfer.

## Methods

### Participants

Thirty-six participants took part in the experiment, 18 male and 18 female with an average age of 20.77 years (*SD* = 2.40). Exclusion criteria were experience in dancing and known problems with VR environments (e.g. motion sickness). The experiment was approved by the ethics committee of the Faculty of Human Sciences at the University of Bern (approval number: 2024-03-02) and was conducted in accordance with the Declaration of Helsinki. All participants provided written informed consent. People shown in the video of the experimental task also provided written consent for publication.

### Experimental task: Joint walking

In the experiment, participants had the role of the follower and were instructed to coordinate their steps with the leader (see video https://doi.org/10.5281/zenodo.23473136 [37]). The leader was one of seven research assistants (three females, four males, randomly assigned to the participants) extensively trained for the task. The joint walking task is inspired by Argentine tango [36]. The participant and the leader faced each other, with the participant’s right hand on the leader’s chest (see Figure 1a). The leader walked forwards and backwards with a varying number of steps to slow tango songs which were partly stretched or compressed to match a constant tempo of 120 bpm. A metronome click marked every second musical beat to increase musical clarity for the participants. The leader took 60-cm steps at a steady cadence of 60 steps per minute, matching the metronome clicks. Leaders were instructed to maintain this pace and step length independently of the participant’s walking quality. One trial lasted 2:50 minutes and contained 160 steps in total distributed over 10 backwards-forwards sequences. One sequence included a fixed number of leader’s forward steps before a change of direction followed by the same number of steps backwards. The change of direction was performed with a closing of the feet and a weight shift from one leg to the other. Across sequences, the number of forward (and thus also backwards) steps varied from five to nine, with each number of steps occurring twice in one trial, randomized over all trials and participants. The number was presented to the leader on a small screen not visible to the participant. This created uncertainty about the upcoming steps and thereby prevented preplanning. However, since forward and backward steps always matched in number, participants could anticipate the forward steps once they knew how many backward steps had occurred. We therefore only analysed the backward steps and excluded the first and last step to remove the acceleration and deceleration phases (see Figure 1b). At the beginning of each trial, a starting tone and eight metronome beats without music allowed the leader and the participant to synchronize through small weight shifts from left to right. The task was performed either in the real world or in a VR environment using an HMD (Vive XR Elite, HTC, Taoyuan, Taiwan, see Figure 1c). In the VR condition, the leader was still physically present, and the hand-to-chest contact remained real, though the visual information was presented virtually.

**Figure 1:**
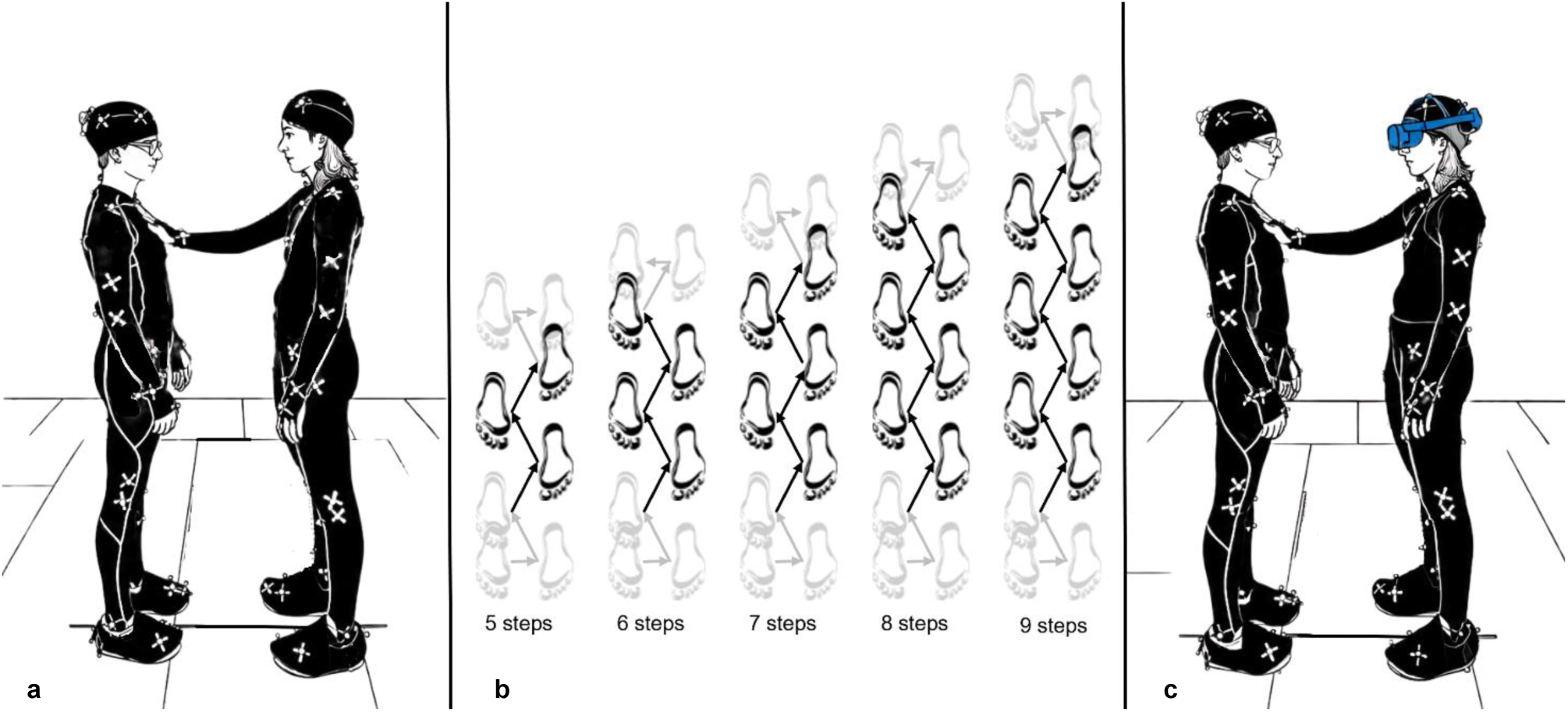
Joint walking task setup and step selection. (a) Leader and participant in the starting position, with the participant’s right hand on the leader’s chest. (b) The different sequences with the five possible numbers of steps before a change of direction varying from five to nine. Black arrows and feet indicate which backward steps were included in the analysis. (c) Both walkers in starting position with the participant wearing the HMD.

To ensure consistency, the leaders’ performance was evaluated after each trial, and they received feedback through an in-house developed software. Trials were aborted and repeated if the leader reported a deviation from the indicated number of steps or the use of the wrong foot. Independently of this, we assessed whether the leader stayed on beat by checking for each backward step whether the velocity minimum and maximum fell within a predefined time window relative to the beat (0.5 s before and after a beat for the maximum; between two beats for the minimum). Trials were repeated if the leader was off beat in more than two steps.

### Motion capture and VR implementation

Full-body motion was tracked using the OptiTrack Motion Capture System (NaturalPoint, Corvallis, USA) comprising eighteen wall-mounted cameras (Prime 13W and Prime 17W) at a sample rate of 200 Hz. Both follower and leader wore whole-body Velcro suits, on which 57 reflective markers with Velcro bases were placed over relevant anatomical landmarks according to the biomechanical full-body marker set from Motive (Motive 2.1.1 NaturalPoint, Corvallis, USA). Both walkers were calibrated with the full marker set in a T-pose, standing upright, feet hip-width apart, with both arms abducted to shoulder height and elbows extended. From that pose measurement, Motive calibrated a skeleton model of each person (see Figure 2a). The eight calibration markers specified in the Motive manual were then removed from each person. To synchronize the motion capture data with the music, we used Streamix (https://doc.streamix.philhum.unibe.ch/), an in-house developed software package based on the theoretical work of Maurer [38]. To represent the walkers in the VR environment, the movements of the leader and the participant were streamed in real time from the motion tracking data into the VR environment using the *Live Link Plugin* from OptiTrack in Unreal Engine. Both walkers were visualized as human-like avatars with resources from the OptiTrack plugin (see Figure 2b).

**Figure 2:**
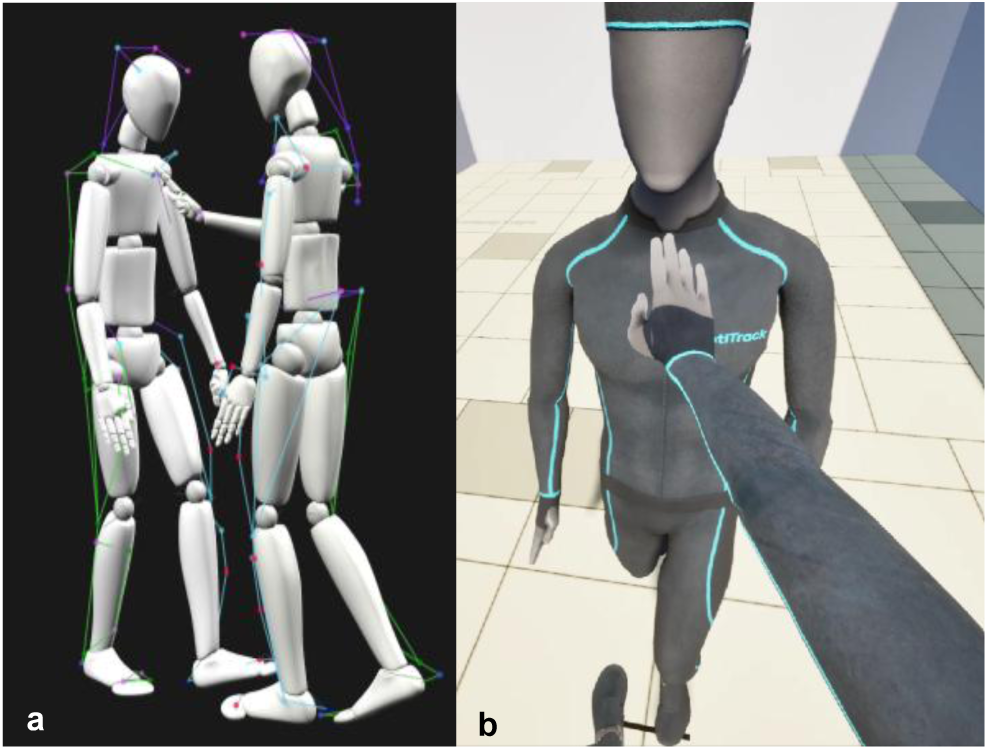
Motion capture setup and VR visualization. (a) Motion-capture representation of the joint walking task. (b) Visualization of the leader as an avatar in the VR environment from the participant’s view.

## Measures

### Dispersion of temporal differences

To quantify leader-follower coordination, we used measures validated and applied by Truffer et al. [36] and calculated them identically for the real and the virtual conditions. We defined the dispersion of temporal differences a priori as our primary dependent variable. It quantifies the stability of the temporal coordination between the leader and follower during one trial (50 backward steps) and was calculated as follows: For each valid backward step, we detected the time points of the leader’s and follower’s peak chest acceleration relative to the beat (Figure 3). We then calculated the temporal difference (Δ*t*, in s) between the two acceleration peaks by subtracting the time point of the follower’s peak from that of the leader’s peak: Δ*t* = t_leader_ − t_follower_. For each trial and participant, we calculated the standard deviation (*SD*) of Δ*t* across all backward steps, yielding the *dispersion of temporal differences, SD(*Δ*t)*. Lower *SD(*Δ*t)* values indicate that the temporal leader-follower coordination is more consistent across steps, reflecting more stable joint walking.

**Figure 3:**
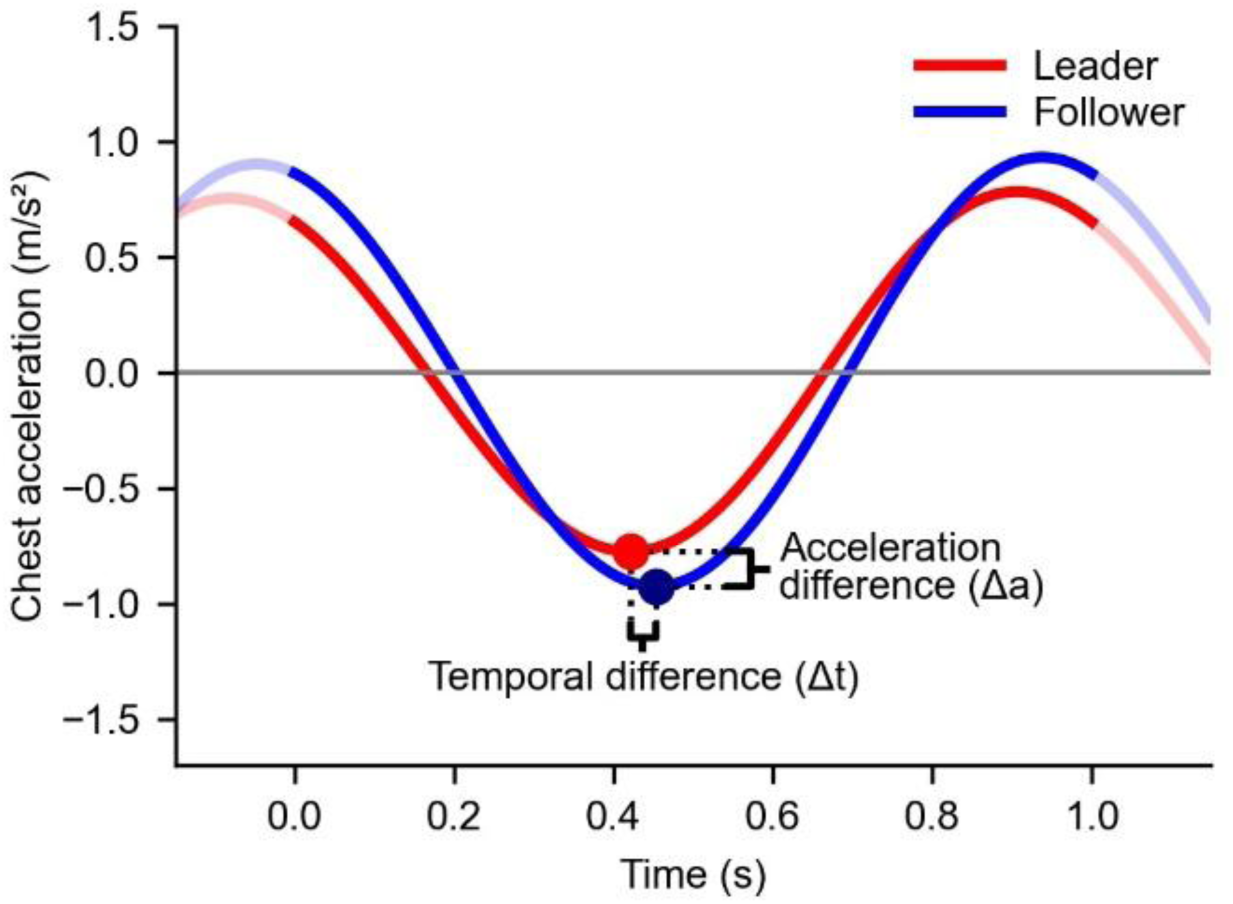
Derivation of the key performance measures. Typical chest acceleration pattern during one step, with beat at 0.0 s and 1.0 s, from which the four dependent variables were derived. Note: since only backward steps were analysed, the maximum accelerations in walking direction are negative peak values.

### Additional leader-follower coordination measures

Following [36], we also calculated three additional coordination measures to provide a more detailed description of the coordination dynamics and to explore whether changes in temporal consistency were accompanied by changes in other aspects of coordination. The first additional measure was the mean of the temporal differences, *M*(Δ*t*). Negative *M(*Δ*t*) values indicate that, on average, the follower reaches the maximum acceleration later than the leader.

Complementing the temporal difference measures, we also assessed the mean and dispersion of absolute acceleration differences between leader and follower, *M(*Δ*a)* and *SD(*Δ*a)*, respectively (see Figure 3), again subtracting the follower’s values from the leader’s (Δ*a =* a_leader_ − a_follower_). Positive *M(*Δ*a)* values indicate a larger peak acceleration for the follower than for the leader and have been interpreted as reflecting a more dynamic, spring-like coordination pattern in which the follower takes up the leader’s input and then accelerates pronouncedly to “catch up” [36]. *SD(*Δ*a)*, in turn, reflects how stable this dynamic coordination pattern is across steps.

### Virtual embodiment

To assess the participant’s subjective perception of the virtual environment, we used an adaptation of the German version of the Virtual Embodiment Questionnaire (VEQ) [39], complemented with questions covering the perceived realism of the virtual leader (see full questionnaire translated to English in Additional file 1, S1). The original questionnaire includes 12 statements about the perception of one’s own body in a virtual environment, divided into the categories *Acceptance/Body Ownership* (e.g. “it felt like the virtual body was my body”), *Control/Agency* (e.g. “the movements of the virtual body felt like they were my movements”), and *Change* (e.g. “I felt like the form or appearance of my own body had changed”). These statements are rated on a seven-level Likert scale from “strongly disagree” (1) to “strongly agree” (7). We added a fourth category *perceived realism of the virtual leader* including six statements in a similar format (e.g. “it felt like the virtual leader was a real person”), as well as one question asking whether it was more difficult to follow the virtual leader than the real leader. As recommended by Roth and Latoschik [39], we computed the mean of all statements within each category.

### Experimental design and procedure

The study consisted of one session lasting three hours. To familiarize participants with the joint walking task, they first walked with the leader forwards and backwards without music and with the number of steps communicated in advance. Participants then had one warm-up trial in each environment (virtual, real) with music. Before the VR trial, participants had time to explore the VR environment and their own movements in the VR world.

The experiment itself consisted of twelve trials: a pre-test with two trials (one in the real, and one in the virtual test environment; order randomized across participants), eight intervention trials, and a post-test with two trials, again one in each environment and in the same order as the pre-test (see Figure 4). After the post-tests, participants completed the adapted VEQ. All pre- and post-tests were performed to the song "El Choclo" (Ángel Villoldo, Orquesta Típica Victor, 1929, public domain). The intervention trials alternated between "El Choclo" and two additional tango songs, "Victoria" (Feliciano Latasa, Orquesta Francisco Canaro, 1935, public domain) and "Condena" (Enrique Santos Discépolo and Francisco Pracánico, Orquesta Francisco Lomuto, 1937, public domain). In all trials, the music was played at a constant volume through a 3^rd^ order Ambisonics sound system rendering two virtual stereo loudspeakers located at about ear-height at the corners of the headwall of the room.

**Figure 4:**
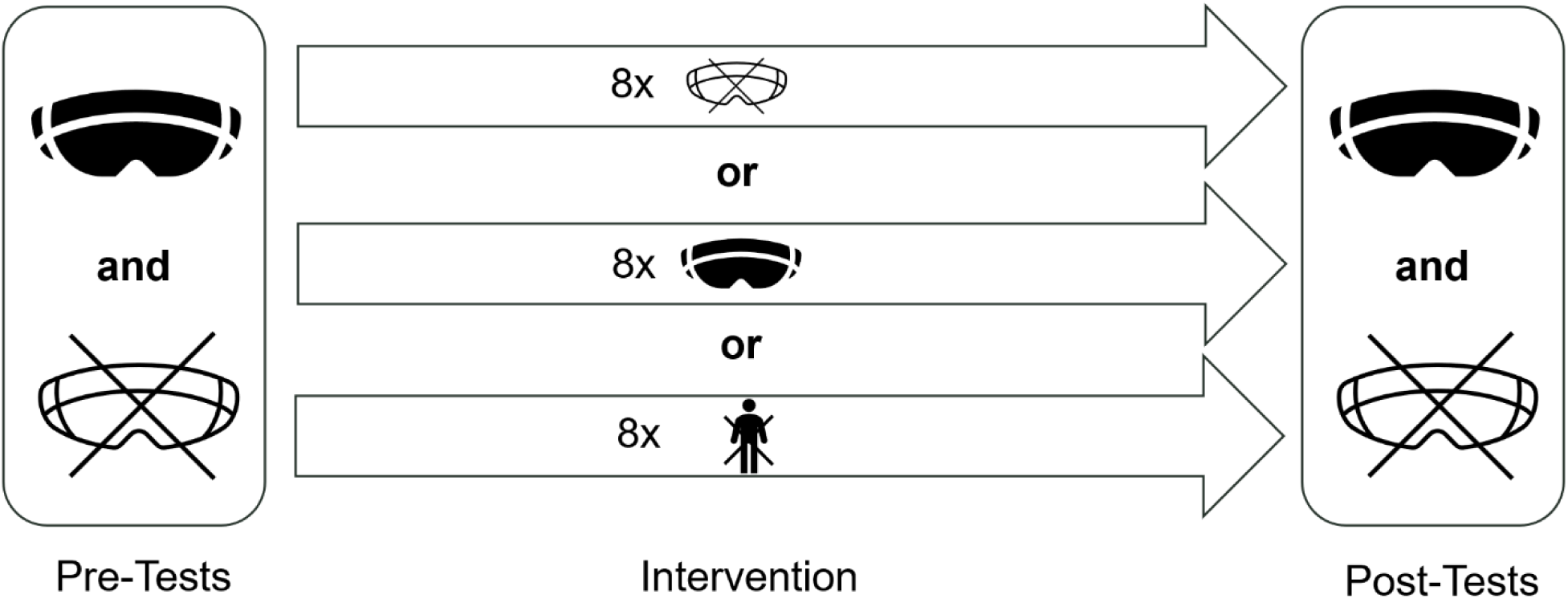
Overview of the study design. Study procedure with one pre-test in each environment (real and virtual), eight intervention trials across three training groups: either training joint walking solely in the real world without the use of VR, solely in the virtual environment using the HMD or walking solo with the leader sitting in front and counting the numbers, followed by one post-test again in each environment.

For the eight intervention trials, participants were divided randomly into three training groups (*n* = 12 per group): *real, virtual*, and *solo*. Groups were randomly assigned before the pre-test, though with sexes equally distributed across groups. In the real training group, participants performed all intervention trials in the real environment, training the joint walking task with no use of VR. In contrast, in the virtual joint walking training group, all intervention trials were performed with the leader physically present and visually displayed in the VR environment using the HMD. In the solo training group, instead of walking together with the leader, the participants performed the intervention trials alone with the leader sitting in front of them and counting the steps aloud. This served as a control group without interpersonal coordination. After each training trial, participants received feedback on their performance with the goal of improving it. In the real and the virtual training group, this value was calculated in the same way as *SD(*Δ*t)* described above but using the velocity instead of the acceleration. In the solo training group, as there was no leader, the *SD* of the time difference between the participant’s velocity peak and the beat onset was calculated instead. Participants were instructed to reduce this value as much as possible.

### Data pre-processing and cleaning

Before aggregating our measures for the pre- and post-test, we removed invalid steps based on the chest velocity, chest acceleration and foot velocity. The chest segment was calculated from the mean position data of the seven thorax and shoulder markers of each participant and their respective leader (Motive convention: SJN, SXS, CV7, TV2, TV7, LHGT, RHGT). Chest velocity and acceleration curves were calculated and filtered with a Savitzky-Golay filter from SciPy version 1.14 (Window Length = 0.9 s, Polyorder = 3) [40]. The curves were then cut according to the metronome beats to obtain data for each step. As described above, only the backward steps without the first and last step of each sequence were considered for data cleaning and analysis. This resulted in 50 steps per participant per trial. Due to recording issues, one pre-test of one participant only included 40 instead of 50 steps, and one post-test of one participant only included 20 instead of 50 steps. This resulted in a total of 7160 steps across all trials.

Steps were counted as valid if the participant and leader performed the backward step with the correct foot, as determined by foot-marker velocities, and if extreme values for velocity and acceleration in the backward direction could be detected within a 1 s interval. This interval began either 0.5 s before a beat for maxima or on a beat for minima. Based on these criteria, we excluded 13 steps (3 in the pre-test data, 10 in the post-test data), leaving 7147 steps.

The remaining data were checked for outliers on a trial-by-trial basis. Steps were labelled as outliers based on the dispersion of velocity extreme values, as well as differences in velocity extreme values between leader and participant, both in magnitude and timing. We excluded steps in which at least one of these measures exceeded three standard deviations within the respective trial. Outliers were distributed similarly across the real (pre-test: 3.22%, post-test: 2.60%) and virtual environments (pre-test: 2.91%, post-test: 3.30%). After data cleaning, 3478 valid steps remained for the pre-tests and 3454 valid steps for the post-tests. Across trials, the median number of valid steps was 48.50 for the pre-tests (*SD* = 1.60, range = 38–50) and 48.00 for the post-tests (*SD* = 3.70, range = 20–50).

### Statistical analysis

To test changes from pre- to post-test and whether these changes differ between groups (virtual, real, and solo), we fitted separate linear mixed-effects models (LMM) for our primary dependent variable, the dispersion of temporal differences, *SD(*Δ*t),* in the real and virtual test environments. The same analysis procedure was applied to the further coordination variables, *M*(Δ*t*), *M(*Δ*a),* and *SD(*Δ*a).* For each participant, two measurements per dependent variable were available in each environment (pre-test and post-test), resulting in 72 observations from 36 participants per environment. We built the LMMs in a sequential, consecutive manner [41] with the *nlme* package [42] (Version 3.1.168) in R (Version 4.5.3) using RStudio (Version 2026.04.0). We set the alpha level at 5%. Fixed-effect coefficients were tested using Wald-type t-tests. Degrees of freedom were estimated with the within-group containment method as implemented in *nlme* [43].

For each variable and environment, model building started from a baseline model with no random effects (grand mean, estimated via generalized least squares) and proceeded stepwise: a random intercept for participant was added first, followed by the fixed effect of time (pre-test vs. post-test). Group was subsequently added, followed by the interaction between time and group. Each model was compared to the previous, less complex model using the log-likelihood ratio *χ^2^* test, as well as the Akaike information criterion (AIC) and the Bayesian information criterion (BIC). For both criteria (AIC, BIC), lower values indicate better model fit, and the magnitude of the decrease (Δ*AIC* and Δ*BIC*) was used to assess the relative support for the improved model [44, 45]. We used the BIC as our main decision criterion due to its stronger penalty for model complexity, consistent with the principle of parsimony. Models were fitted using maximum likelihood for model comparisons; the final model was refitted using restricted maximum likelihood (REML) for parameter estimation. The best-fitting, most parsimonious model was retained as the final model for statistical inferences for each variable and environment.

To rule out that the observed effects depended on the order in which time and group were entered for sequential testing, we checked the reverse predictor order (group before time) for all four coordination variables in both environments. For all variable-environment combinations, the interpretations of the resulting model were identical to the initial order.

As the dependent variables already contained aggregated data per trial (mean and standard deviation across up to 50 steps), there were no missing values at this stage. Normality and homoscedasticity were graphically checked (diagnostic plots are available in the repository [37]). Due to skewness in the *SD(*Δ*t)* residuals, both final models were re-estimated using a case bootstrap at the participant level (i.e. resampling participants with their pre- and post-test observations; 5,000 iterations) in R (lme4, lmerTest), with 95% confidence intervals (CIs) derived from the 2.5th/97.5th percentiles of the bootstrap distributions. Bootstrap estimates and CIs were consistent with the original REML models, confirming the robustness of the reported effects. At the observation level (level 1), no outliers were found based on Cook’s distance with a threshold set to 1 [41]. At the participant level (level 2), potential outliers were identified by extracting the random intercepts and flagging participants with an absolute z-score greater than 2, resulting in no more than three extreme participants per model. Because sensitivity analyses showed that excluding these individuals did not change the interpretation of the results, all participants were retained in the final analysis. To assess potential multicollinearity among the independent variables where applicable, variance inflation factors (VIF) were examined and found to be close to 1.0, well below the standard conservative threshold of 5.0 [46], indicating that multicollinearity was not an issue. The data and analysis code required to reproduce the reported statistical analyses are available in the accompanying Zenodo repository [37].

## Results

### Dispersion of temporal differences

Figure 5 shows model-estimated means in the dispersion of temporal differences, *SD(*Δ*t)*, before and after training in the real (left) and virtual test environment (right). The model-estimated means are derived from the full model including time, group, and the time × group interaction.

**Figure 5:**
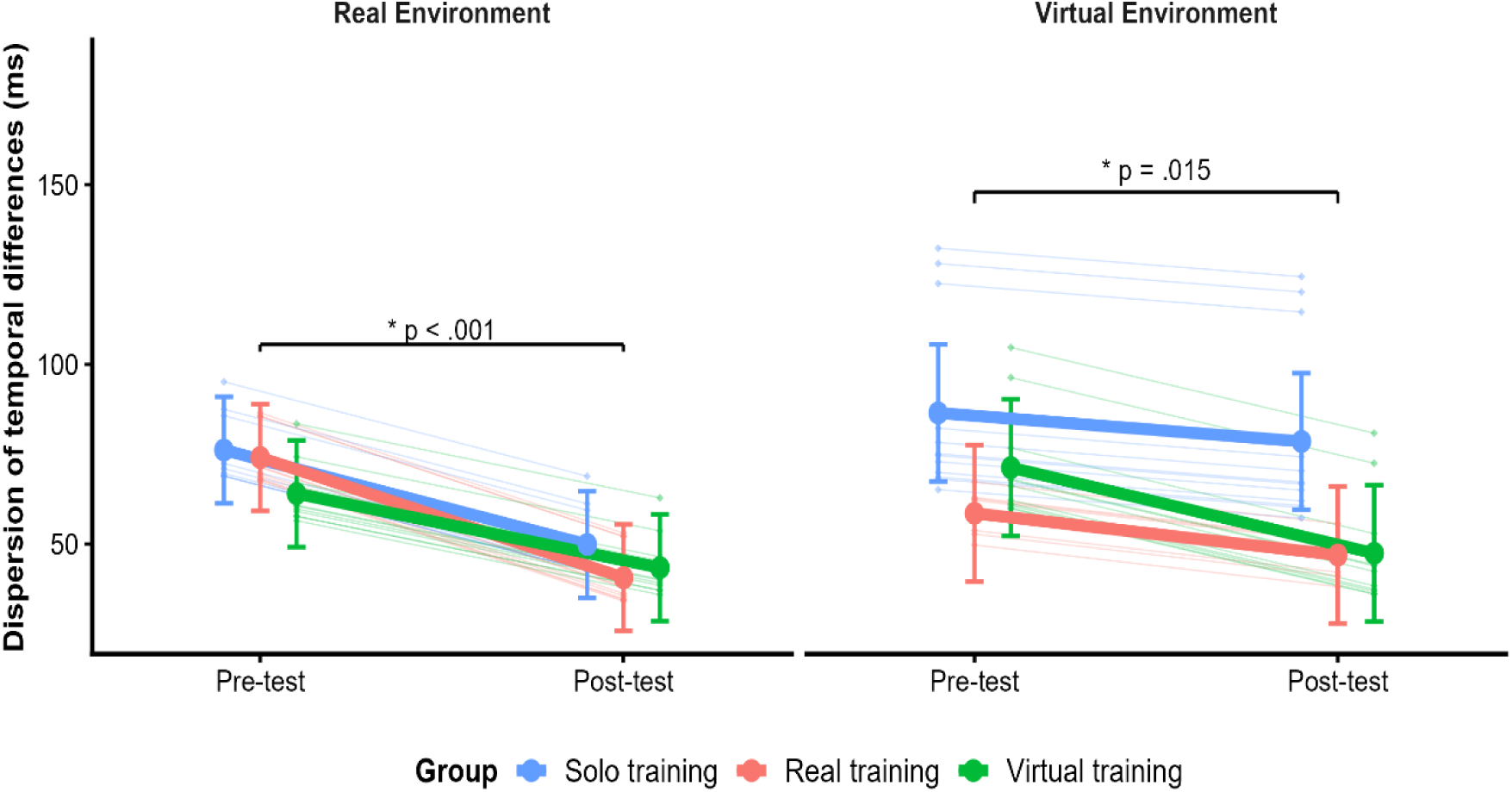
Dispersion of temporal differences, SD(Δt), before and after training in both test environments. Thin lines show model-fitted individual trajectories per participant; bold lines and error bars show model-estimated marginal means with 95% confidence intervals, separately for the three training groups. Brackets show the pre- to-post contrast averaged across training groups with its p-value. All values are derived from the full model including time, group, and their interaction (random intercept for participant). The statistically retained final model for SD(Δt) included only the fixed effect of time (see Results); groups are displayed for descriptive purposes only (n = 36 participants, 72 observations per environment).

Participants who trained in VR showed pre-post improvements (i.e. reductions in *SD*[Δ*t*]) in both the virtual test environment (*M*_pre_ = 71.27, 95% CI_pre_ = [52.25, 90.28]; *M*_post_ = 47.42, 95% CI_post_ = [28.40, 66.44]) and the real test environment (*M*_pre_ = 64.01, CI_pre_ = [49.18, 78.83]; *M*_post_ = 43.40, CI_post_ = [28.57, 58.22]). A similar cross-environment pattern was found in the real training group, which also showed improvements in both the real (*M*_pre_ = 74.06, CI_pre_ = [59.24, 88.89]; *M*_post_ = 40.61, CI_post_ = [25.79, 55.44]) and the virtual test environment (*M*_pre_ = 58.54, CI_pre_ = [39.52, 77.56]; *M*_post_ = 46.93, CI_post_ = [27.91, 65.95]). These improvements in the joint walking training groups were, however, not significantly greater than those observed in the solo walking group (real environment: *M*_pre_ = 76.15, CI_pre_ = [61.33, 90.98]; *M*_post_ = 49.87, CI_post_ = [35.05, 64.69]; virtual environment: *M*_pre_ = 86.47, CI_pre_ = [67.45, 105.49]; *M*_post_ = 78.55, CI_post_ = [59.53, 97.57]).

For both test environments, models including the time × group interaction did not provide a better fit than the models without the interaction (see Additional file 2: Tables S1–S4). For the real test environment, the model including only the fixed effect of time and a random intercept for participant was retained (see Additional file 2: Table S1). This model (*R*^2^_marginal_ = .23, intraclass correlation coefficient [ICC] = .25) showed a significant improvement from pre- to post-test across groups (*b* = −26.78 [−37.09, −16.47], *t*(35) = −5.27, *p* < .001, see Additional file 2: Table S2), indicating that leader–follower coordination became more consistent from pre- to post-test irrespective of training group. For the virtual test environment, also the model including only the fixed effect of time and a random intercept for participant was retained (see Additional file 2: Table S3). Similarly, this model (*R*^2^_marginal_ = .04, ICC = .51) showed general improvements across groups (*b* = −14.46 [−25.85, −3.06], *t*(35) = −2.58, *p* = .014, see Additional file 2: Table S4).

### Further coordination variables

Figure 6 shows model-estimated marginal means of *M(*Δ*t)*, *M(*Δ*a)* and *SD(*Δ*a)* before and after training in both test environments (see also Additional file 2: Table S5). For *M(*Δ*t)*, the model with only a random intercept for participants was retained in both environments (see Additional file 2: Tables S6–S9). *M(*Δ*t)* differed significantly from zero in the real environment (*b* = 15.75 [2.03, 29.47], *t*(36) = 2.33, *p* = .026, ICC = .76), but not in the virtual environment (*b* = 3.02 [-10.41, 16.46], *t*(36) = 0.46, *p* = .651, ICC = .68).

**Figure 6:**
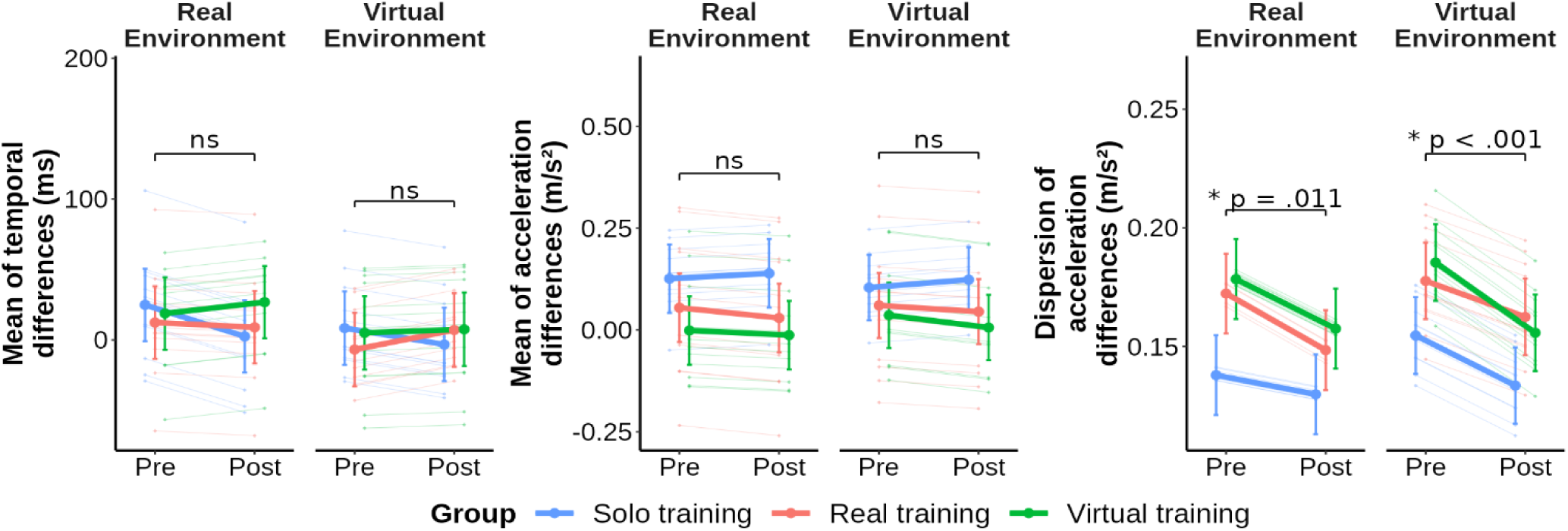
Development of M(Δt), M(Δa) and SD(Δa) from pre- to post-test in both test environments. Thin lines show model-fitted individual trajectories per participant; bold lines and error bars show model-estimated marginal means with 95% confidence intervals, separately for the three training groups. Brackets show the pre- to-post contrast averaged across training groups with its p-value (ns = not significant). All values are derived from the full model including time, group, and their interaction (random intercept for participant) and are shown for visualization purposes (see Results for the statistically retained models; n = 36 participants, 72 observations per environment).

Also, for *M(*Δ*a)*, the model with only a random intercept for participants was retained in both environments (see Additional file 2: Tables S10 and S11 for the real and S12 and S13 for the virtual environment). In these models, *M(*Δ*a)* was significantly above zero in both the real (*b* = 0.06 [0.01, 0.11], *t*(36) = 2.28, *p* = .029, ICC = .89) and the virtual environment (*b* = 0.06 [0.02, 0.11], *t*(36) = 2.80, *p* = .008, ICC = .87). For *SD(*Δ*a)*, the model including time and group as fixed effects and a random intercept for participants was retained in both environments (see Additional file 2: Tables S14 and S15 for the real and Tables S16 and S17 for the virtual environment). In these models (real: *R*^2^_marginal_ = .27, ICC = .07, virtual: *R*^2^_marginal_ = .27, ICC = .54), *SD(*Δ*a)* decreased significantly from pre- to post-test (real: *b* = -0.02 [-0.03, -0.00], *t*(35) = -2.72, *p* = .010, virtual: *b* = -0.02 [-0.03, -0.01], *t*(35) = -5.00, *p* < .001). The real training group (real: *b* = 0.03 [0.01, 0.04], *t*(33) = 3.11, *p* = .004, virtual: *b* = 0.03 [0.01, 0.05], *t*(33) = 2.64, *p* = .012) and the virtual training group (real: *b* = 0.03 [0.02, 0.05], *t*(33) = 4.00, *p* < .001, virtual: *b* = 0.03 [0.01, 0.05], *t*(33) = 2.70, *p* = .011) showed significantly higher values than the solo training group in both environments.

Overall, the two types of coordination measures showed the same pattern. For both the temporal difference and the acceleration difference measures, the *SD* decreased from pre- to post-test, while *M* did not change. The group difference in *SD(*Δ*a)* did not interact with time and therefore likely reflects a stable baseline difference rather than a practice-related improvement.

As a robustness check, we modelled all four coordination variables in a combined dataset including both test environments (see Additional file 2: Table S18). The combined analysis yielded the same final models as the separate analyses (BIC-based selection), and adding environment to the retained models did not improve model fit for *SD(*Δ*t)*, *SD(*Δ*a)*, and *M(*Δ*a)*. For *M(*Δ*t)*, adding environment to the retained model improved model fit (*χ*²(1) = 10.07, *p* = .002, Δ*BIC* = −5.1).

### Virtual embodiment questionnaire

To assess participants’ subjective experience of the virtual environment, all participants completed the VEQ. Mean subscale scores were close to the scale midpoint for the category Acceptance/Body Ownership (*M* = 4.20, *SD* = 1.17) and slightly below it for the category Change of one’s own body in the virtual world (*M* = 3.63, *SD* = 1.28). Scores were moderately high for the category Control/Agency over the virtual body (*M* = 5.28, *SD* = 0.78), indicating that participants experienced a sense of control over the virtual body’s movements and perceived them as synchronized with their own. The category Perceived realism of the virtual leader was moderate (*M* = 4.50, *SD* = 1.00), while participants reported that following the virtual leader was more difficult than following the real leader (*M* = 5.83, *SD* = 1.42).

## Discussion

In this study, we investigated the effects of joint walking training in virtual and real environments and the transfer of training-related improvements between environments. Participants trained either with a real partner, with the partner displayed as an avatar in VR or walked to the beat without a partner. The results show that participants who trained in VR improved the consistency of leader-follower coordination in both the virtual and the real test environments, consistent with VR-to-real-world transfer. Similarly, participants who trained in the real-world environment showed improvements in the virtual test environment. However, improvements in the leader-follower coordination were also observed in participants who practised walking to the beat without a partner.

### Transfer from virtual training to the real-world environment

With the increasing use of VR for training across domains – including rehabilitation, sports, and medical education – understanding whether improvements achieved in virtual environments transfer to real-world performance is becoming increasingly important [17, 23, 28, 47]. Our results show that participants who practised joint walking in VR improved not only in the virtual but also in the real-world test environment. The magnitude of these improvements was comparable to that observed in participants who trained in the real-world condition.

These findings are consistent with previous work demonstrating positive transfer of VR training to real-world performance across different motor tasks (e.g. obstacle crossing during walking [23]; sensorimotor assembly [48]; golf putting [49]; table tennis [50–52]; karate technique [16]; baseball batting [25]; robot-assisted surgery [47]; post-stroke rehabilitation of gait parameters [53] and arm reaching [54]; obstacle negotiation in older adults and people with Parkinson’s disease [32]). Our findings extend this empirical basis to interpersonal coordination during walking. However, there are also studies reporting no transfer after VR training (e.g. response behaviour in karate [55]; balance after stroke [56]) or even negative transfer (e.g. dart throwing [57]; laparoscopic surgery [58]; isometric pinch force task [59]). These mixed findings suggest that successful transfer depends on whether the VR task captures functionally relevant components of the real-world task and enables learning its functional relationships. This aligns with structural learning theories, which propose that learners extract invariant features shared by related tasks, facilitating transfer to new tasks with the same structure [60] (for VR-training transfer, see [61]). When task-relevant features are preserved, physical parameters may be modified and the visualization simplified without preventing transfer [27]. Reviews in rehabilitation [17] and sports [62] similarly emphasized that VR does not need to fully replicate reality (e.g. in visual appearance), but that the fidelity of task-relevant sensorimotor interactions is likely to be important for transfer.

Our task elicited similar behaviour in the virtual and real environment. Comparing the different measures of leader-follower coordination between the virtual and real environments (Figure 5, Figure 6) indicates their movement dynamics and strategies did not systematically differ between the environments. This is in line with participants’ subjective experiences, with participants reporting a relatively high sense of agency and controllability over their virtual body, while perceiving the virtual leader as moderately realistic and reporting higher difficulty following the virtual than the real leader. Thus, while the virtual environment was not perceived as fully equivalent to the real environment, it felt controllable to the participants and elicited comparable behaviour.

### Contributions of joint and solo walking

While our results support the hypothesis that training interpersonal coordination in VR extends to the real world, we observed that participants who practised rhythmic walking to the beat showed comparable improvements in the joint walking task. The absence of a time × group interaction prevents us from disentangling a specific transfer of the acquired interpersonal coordination skill in VR to the real-world from a more general practice effect.

Nevertheless, the improvements of the solo group may provide insights into task-relevant dimensions and potentially transferable components in the task. One possible interpretation is that performance improvements in our task (i.e. reducing dispersion of temporal differences in leader-follower coordination, *SD[*Δ*t]*) may be driven by two processes: The first one is improving the use of the leader’s multisensory (visual and haptic) signals in the continuous control of action. This aspect is practised in the joint walking but not in the solo walking training. Second, because the leader also synchronizes to the same audio beat, improving consistent stepping relative to the beat may also improve performance in the interpersonal task. The solo walking group specifically trained walking to the beat for the same number of trials and to the same music. Hence, improvements may be driven by this second approach, especially in the solo group. That participants use auditory information from the beat in this task was shown by Truffer et al. [36], who found that removing it significantly increased *SD*(Δ*t)*. This effect was, however, substantially smaller than the effect found when haptic information was removed, suggesting that auditory cues alone are unlikely to fully account for coordination performance and that online adjustments to the partner remain important. The potential contribution of individual rhythmic abilities is also supported by findings from dyadic finger tapping. Pecenka and Keller [63] found that the variability of individuals synchronizing with a pacing signal explained about one third of the variability of synchronization in dyads.

To disentangle these contributions, future studies could increase the demands for real-time partner-based coordination by introducing higher uncertainty in the leader’s movements. Although the number of consecutive steps was unknown to the participants in the current study, it is likely that a consistent coordination to the beat drives a large part of the performance. Introducing variable step lengths or lateral steps could increase the need for partner-specific movement information that cannot be derived from the beat alone. Such manipulations may help determine whether joint walking training provides additional benefits beyond individual rhythmic stepping practice. Examining whether – and under which conditions or populations – joint walking offers additional (or complementary) benefits compared to solo walking [10] is also relevant for rehabilitation, given the wide use of rhythmic auditory cueing to support gait and motor rehabilitation [64–66].

### Potential for rehabilitation

Overall, our findings provide initial support for the potential of VR-based joint walking interventions in older adults or clinical populations. We found no indications that training in VR was less effective in improving joint walking coordination than training in the real-world environment. Additionally, we found that joint walking in VR elicited behaviour similar to that observed in the real-world condition and maintained a subjective feeling of agency. Although these findings are encouraging, further research is needed to establish the clinical relevance of VR-based joint walking training. Beyond disentangling the mechanisms of joint and solo walking to a beat, future studies should investigate longer training interventions in older adults and clinical populations and assess whether improvements persist over time. The present study provides initial evidence consistent with “near transfer” [67, 68], as improvements following VR training extended to the same task performed in the real-world environment. For rehabilitation applications, an important next step is to investigate “far transfer” effects, that is, whether these improvements generalize beyond the trained task to clinically relevant outcomes, such as gait speed, cadence, step length, and fall risk.

Solo walking to a beat (i.e. rhythmic auditory cueing) has been shown to improve gait velocity, stride length and cadence in younger and older adults [69], and our results indicate that it can also improve the consistency of leader–follower coordination in our joint walking task. However, dance-related research suggests that moving with a partner may offer additional benefits. A meta-analysis reported positive effects of tango dancing on motor severity, balance and the Timed Up and Go test in people with Parkinson’s disease [70]. In older adults at risk of falls, tango tended to improve balance and walking speed more than a walking programme [71]. Hackney and Earhart [72] attributed such effects not only to the music but also to physical cues by partners. The partner’s weight shifts and direction of movement can provide cues that help to initiate steps and to maintain stride length and cadence. Moreover, partner dance such as tango involves improvised step patterns, so that the follower must continuously adjust to the partner’s movements [70, 72]. This capability to adapt gait to unforeseen changes is often reduced in older adults and may increase their risk of falls [3]. Our task includes this continuous adaptation to the leader’s movements under uncertainty in a simplified form.

An important consideration for future applications is that VR training in the present study combined virtual and physical interaction. For practical applications, training with a virtual partner without requiring another person would be of great practical value. However, haptics plays a major role in interpersonal coordination [36, 73, 74]. Therefore, two complementary approaches could be explored to enable training without a human partner: First, it has been shown that synchronization to partners based on visual and auditory cues can shape gait [34, 35]. For example, when walking behind a leader, followers adapted their stride-time dynamics to those of the leader [7]. How the present results would change without the presence of a physical partner, and whether such training would be accepted by users, needs to be tested in future studies. Second, virtual partners could be combined with robotic devices to provide physical cues without requiring a human partner. Research has shown that older adults accept robots as partners, both as a walking companion [75] and in a forward and backward stepping task where they led the robot through physical contact [76]. Combining an immersive virtual partner with robotic haptic assistance could therefore provide a promising direction for developing partner-independent training systems. In this context, our study provides a first step towards a VR-based joint walking training by establishing a baseline understanding of learning and transfer effects, suggesting that it may offer a flexible and scalable approach for gait rehabilitation.

## Conclusions

Joint walking training in VR resulted in improved leader–follower coordination in both a virtual and a real-world test environment, consistent with a VR-to-real-world transfer effect. Participants trained in VR showed improvements comparable to those observed after real-world training. These findings provide initial support for the potential of VR-based joint walking training as a promising approach for gait training and rehabilitation. However, comparable improvements following solo walking suggest that improved consistency of stepping relative to the auditory beat may have contributed to the observed improvements in interpersonal coordination. Future studies should therefore clarify the specific contribution of partner-based interaction and examine whether such training improves clinically relevant gait outcomes.

## List of abbreviations

AIC: Akaike information criterion
BIC: Bayesian information criterion
CI: Confidence interval
HMD: Head-mounted display
ICC: Intraclass correlation coefficient
LMM: Linear mixed-effects model
M(Δa): Mean of acceleration differences
M(Δt): Mean of temporal differences
REML: Restricted maximum likelihood
SD: Standard deviation
SD(Δa): Dispersion of acceleration differences
SD(Δt): Dispersion of temporal differences
VEQ: Virtual Embodiment Questionnaire
VIF: Variance inflation factor
VR: Virtual reality

## Declarations

### Ethics approval and consent to participate

The study was approved by the Ethics Committee of the Faculty of Human Sciences at the University of Bern (approval number 2024-03-02) and was conducted in accordance with the Declaration of Helsinki. All participants provided written informed consent prior to participation.

### Consent for publication

Written informed consent for publication was obtained from all individuals identifiable in the accompanying video of the experimental task.

### Availability of data and materials

The datasets generated and analysed during the current study and the analysis code are available on Zenodo, https://doi.org/10.5281/zenodo.23473136 [37]. The repository also contains the diagnostic plots of the model assumptions and a video demonstrating the experimental task.

### Authors’ contributions

**CD:** Conceptualization, Investigation, Data Curation, Formal analysis, Visualization, Writing - Original Draft.

**MT:** Conceptualization, Methodology, Validation, Investigation, Data Curation, Writing - Review C Editing.

**DB:** Conceptualization, Methodology, Software, Formal analysis, Visualization.

**RK:** Funding Acquisition, Conceptualization, Methodology, Software, Writing - Review C Editing.

**SZ:** Funding Acquisition, Project administration, Supervision, Conceptualization, Methodology, Writing - Review C Editing.

Constanze Dammeyer and Mathilde Truffer contributed equally to this work. All authors read and approved the final manuscript.

## Acknowledgements

We are grateful to Martin Widmer for his technical support with the implementation of the VR task and the development of the data export scripts.

## Additional Data

**File name:** Additional file 1

**File format:** .pdf

**Title of data:** English version of the adapted virtual embodiment questionnaire

**Description of data:** English version of the questionnaire on the perception of the virtual environment, including the items of the Virtual Embodiment Questionnaire and the additional items on the perceived realism of the virtual leader.

**File name:** Additional file 2

**File format:** .pdf

**Title of data:** Supplementary Tables S1–S18

**Description of data:** Sequential model comparisons and coefficient tables of the retained and the full interaction model for all four coordination variables, *SD(*Δ*t), M(*Δ*t), M(*Δ*a)* and *SD(*Δ*a)*, in the real and the virtual test environment (Tables S1–S4 and S6–S17). Model-estimated marginal means of *M(*Δ*t), M(*Δ*a)* and *SD(*Δ*a)* for each training group, test environment and time point (Table S5). Sequential model comparison for the combined dataset of both test environments (Table S18).

