## Additional file 2 for "Joint Walking Training with Virtual and Real Partners: Testing the Transfer from Virtual Reality to the Real World"

***Table S1 | Sequential model building and comparison for dispersion of temporal differences, SD(Δt) (real test environment)***

| **Model** | ***df*** | ***AIC*** | ***BIC*** | ***logLik*** | **Test** | **$\chi^2$** | ***p*** | **Δ Marginal $R^2$** |
|---|---|---|---|---|---|---|---|---|
| Model 0: Intercept-only (GLS) | 2 | 688.23 | 692.79 | -342.12 | – | – | – | – |
| Model 1: Random intercept | 3 | 690.23 | 697.06 | -342.12 | 0 vs 1 | 0.00 | p = 1.000 | – |
| Model 2: + Time | 4 | 671.14 | 680.25 | -331.57 | 1 vs 2 | 21.09 | p < .001 | .23 |
| Model 3: + Group | 6 | 673.72 | 687.38 | -330.86 | 2 vs 3 | 1.42 | p = .492 | .02 |
| Model 4: + Time × Group | 8 | 676.60 | 694.82 | -330.30 | 3 vs 4 | 1.12 | p = .572 | .01 |

Note. *GLS* Generalized least squares, *df* Number of estimated parameters, *AIC* Akaike information criterion, *BIC* Bayesian information criterion, *logLik* Log-likelihood, $\chi^2$ Likelihood ratio test statistic against the previous model, *p* p-value, *Δ Marginal $R^2$* Change in marginal $R^2$ (variance explained by fixed effects) compared to the previous model. All models were estimated with maximum likelihood. logLik, AIC and BIC refer to the outcome in ms. An en-dash (–) indicates not applicable.

***Table S2 | Multilevel regression of dispersion of temporal differences, SD(Δt), on time and training group in the real test environment***

| | Model 2: + Time | | | | | Model 4: + Time × Group | | | | |
|---|---|---|---|---|---|---|---|---|---|---|
| **Predictor** | ***B*** | ***SE*** | **95% CI** | ***t (df)*** | ***p*** | ***B*** | ***SE*** | **95% CI** | ***t (df)*** | ***p*** |
| Intercept | 71.41 | 4.16 | [62.97, 79.85] | 17.17 (35) | p < .001 | 76.15 | 7.29 | [61.33, 90.98] | 10.45 (33) | p < .001 |
| Time | -26.78 | 5.08 | [-37.09, -16.47] | -5.27 (35) | p < .001 | -26.28 | 8.92 | [-44.43, -8.14] | -2.95 (33) | p = .006 |
| Group Real | – | – | – | – | – | -2.09 | 10.30 | [-23.05, 18.87] | -0.20 (33) | p = .840 |
| Group Virtual | – | – | – | – | – | -12.15 | 10.30 | [-33.11, 8.82] | -1.18 (33) | p = .247 |
| Time × Group Real | – | – | – | – | – | -7.17 | 12.61 | [-32.82, 18.49] | -0.57 (33) | p = .574 |
| Time × Group Virtual | – | – | – | – | – | 5.67 | 12.61 | [-19.99, 31.33] | 0.45 (33) | p = .656 |
| *Random effects* | | | | | | | | | | |
| Residual variance ($\sigma^2$) | 464.09 | – | – | – | – | 477.17 | – | – | – | – |
| Intercept variance ($\tau_{00}$) | 158.66 | – | – | – | – | 159.79 | – | – | – | – |
| ICC | .25 | – | – | – | – | .25 | – | – | – | – |
| Marginal $R^2$ | .23 | – | – | – | – | .24 | – | – | – | – |
| Conditional $R^2$ | .42 | – | – | – | – | .43 | – | – | – | – |
| Δ Marginal $R^2$ | – | – | – | – | – | .02 | – | – | – | – |
| N subjects | 36 | – | – | – | – | 36 | – | – | – | – |

Note. *B* Unstandardized regression coefficient, *SE* Standard error, *CI* Confidence interval, *t*(df) t-statistic with degrees of freedom (within-group containment method as implemented in nlme::lme; Pinheiro & Bates, 2000), *p* p-value, $\sigma^2$ Residual variance (within-subject variance), $\tau_{00}$ Random intercept variance (between-subject variance), *ICC* Intraclass correlation coefficient, *Marginal $R^2$* Variance explained by fixed effects, *Conditional $R^2$* Variance explained by fixed and random effects, *Δ Marginal $R^2$* Change in marginal $R^2$ from Model 2 to Model 4. Both models were estimated with restricted maximum likelihood (REML). Reference categories: Pre-Test for time and Solo for group. Estimates are given in ms, variances in ms². Model statistics: N_participants = 36, N_observations = 72. An en-dash (–) indicates that a value is not applicable.

***Table S3 | Sequential model building and comparison for dispersion of temporal differences, SD(Δt) (virtual test environment)***

| **Model** | ***df*** | ***AIC*** | ***BIC*** | ***logLik*** | **Test** | $\chi^2$ | ***p*** | **Δ Marginal $R^2$** |
|---|---|---|---|---|---|---|---|---|
| Model 0: Intercept-only (GLS) | 2 | 718.10 | 722.65 | -357.05 | – | – | – | – |
| Model 1: Random intercept | 3 | 712.03 | 718.86 | -353.01 | 0 vs 1 | 8.07 | p = .005 | – |
| Model 2: + Time | 4 | 707.78 | 716.88 | -349.89 | 1 vs 2 | 6.25 | p = .012 | .04 |
| Model 3: + Group | 6 | 704.22 | 717.88 | -346.11 | 2 vs 3 | 7.56 | p = .023 | .14 |
| Model 4: + Time × Group | 8 | 706.67 | 724.88 | -345.33 | 3 vs 4 | 1.55 | p = .462 | .01 |

Note. *GLS* Generalized least squares, *df* Number of estimated parameters, *AIC* Akaike information criterion, *BIC* Bayesian information criterion, *logLik* Log-likelihood, $\chi^2$ Likelihood ratio test statistic against the previous model, *p* p-value, *Δ Marginal $R^2$* Change in marginal $R^2$ (variance explained by fixed effects) compared to the previous model. All models were estimated with maximum likelihood. logLik, AIC and BIC refer to the outcome in ms. An en-dash (–) indicates not applicable.

***Table S4 | Multilevel regression of dispersion of temporal differences, SD(Δt), on time and training group in the virtual test environment***

| | Model 2: + Time | | | | | Model 4: + Time × Group | | | | |
|---|---|---|---|---|---|---|---|---|---|---|
| **Predictor** | ***B*** | ***SE*** | **95% CI** | ***t (df)*** | ***p*** | ***B*** | ***SE*** | **95% CI** | ***t (df)*** | ***p*** |
| Intercept | 72.09 | 5.70 | [60.53, 83.66] | 12.66 (35) | p < .001 | 86.47 | 9.35 | [67.45, 105.49] | 9.25 (33) | p < .001 |
| Time | -14.46 | 5.61 | [-25.85, -3.06] | -2.58 (35) | p = .014 | -7.92 | 9.80 | [-27.86, 12.01] | -0.81 (33) | p = .425 |
| Group Real | – | – | – | – | – | -27.93 | 13.22 | [-54.83, -1.03] | -2.11 (33) | p = .042 |
| Group Virtual | – | – | – | – | – | -15.21 | 13.22 | [-42.10, 11.69] | -1.15 (33) | p = .258 |
| Time × Group Real | – | – | – | – | – | -3.69 | 13.86 | [-31.88, 24.51] | -0.27 (33) | p = .792 |
| Time × Group Virtual | – | – | – | – | – | -15.92 | 13.86 | [-44.12, 12.27] | -1.15 (33) | p = .259 |
| *Random effects* | | | | | | | | | | |
| Residual variance ($\sigma^2$) | 566.97 | – | – | – | – | 576.07 | – | – | – | – |
| Intercept variance ($\tau_{00}$) | 601.33 | – | – | – | – | 472.62 | – | – | – | – |
| ICC | .51 | – | – | – | – | .45 | – | – | – | – |
| Marginal $R^2$ | .04 | – | – | – | – | .18 | – | – | – | – |
| Conditional $R^2$ | .54 | – | – | – | – | .55 | – | – | – | – |
| Δ Marginal $R^2$ | – | – | – | – | – | .14 | – | – | – | – |
| N subjects | 36 | – | – | – | – | 36 | – | – | – | – |

Note. *B* Unstandardized regression coefficient, *SE* Standard error, *CI* Confidence interval, *t*(df) t-statistic with degrees of freedom (within-group containment method as implemented in nlme::lme; Pinheiro & Bates, 2000), *p* p-value, $\sigma^2$ Residual variance (within-subject variance), $\tau_{00}$ Random intercept variance (between-subject variance), *ICC* Intraclass correlation coefficient, *Marginal $R^2$* Variance explained by fixed effects, *Conditional $R^2$* Variance explained by fixed and random effects, *Δ Marginal $R^2$* Change in marginal $R^2$ from Model 2 to Model 4. Both models were estimated with restricted maximum likelihood (REML). Reference categories: Pre-Test for time and Solo for group. Estimates are given in ms, variances in ms². Model statistics: N_participants = 36, N_observations = 72. An en-dash (–) indicates that a value is not applicable.

***Table S5 | Model-estimated marginal means of M(Δt), M(Δa) and SD(Δa) by test environment, training group and time point***

| | | ***Pre-Test*** | | ***Post-Test*** | |
|---|---|---|---|---|---|
| **Test environment** | **Training group** | ***M*** | **95% CI** | ***M*** | **95% CI** |
| *Mean of temporal differences, M(Δt), in ms* | | | | | |
| Real | Solo | 24.90 | [-0.80, 50.59] | 2.50 | [-23.20, 28.20] |
| Real | Real | 12.38 | [-13.32, 38.08] | 9.02 | [-16.68, 34.72] |
| Real | Virtual | 18.82 | [-6.87, 44.52] | 26.86 | [1.16, 52.56] |
| Virtual | Solo | 8.44 | [-17.63, 34.51] | -3.19 | [-29.25, 22.88] |
| Virtual | Real | -6.80 | [-32.87, 19.26] | 7.06 | [-19.00, 33.13] |
| Virtual | Virtual | 5.08 | [-20.98, 31.15] | 7.55 | [-18.52, 33.61] |
| *Mean of absolute acceleration differences, M(Δa), in m/s²* | | | | | |
| Real | Solo | 0.13 | [0.04, 0.21] | 0.14 | [0.05, 0.22] |
| Real | Real | 0.05 | [-0.03, 0.14] | 0.03 | [-0.05, 0.11] |
| Real | Virtual | -0.00 | [-0.09, 0.08] | -0.01 | [-0.10, 0.07] |
| Virtual | Solo | 0.10 | [0.02, 0.18] | 0.12 | [0.04, 0.20] |
| Virtual | Real | 0.06 | [-0.02, 0.14] | 0.05 | [-0.04, 0.13] |
| Virtual | Virtual | 0.04 | [-0.04, 0.12] | 0.01 | [-0.07, 0.09] |
| *Dispersion of absolute acceleration differences, SD(Δa), in m/s²* | | | | | |
| Real | Solo | 0.14 | [0.12, 0.15] | 0.13 | [0.11, 0.15] |
| Real | Real | 0.17 | [0.16, 0.19] | 0.15 | [0.13, 0.17] |
| Real | Virtual | 0.18 | [0.16, 0.20] | 0.16 | [0.14, 0.17] |
| Virtual | Solo | 0.15 | [0.14, 0.17] | 0.13 | [0.12, 0.15] |
| Virtual | Real | 0.18 | [0.16, 0.19] | 0.16 | [0.15, 0.18] |
| Virtual | Virtual | 0.19 | [0.17, 0.20] | 0.16 | [0.14, 0.17] |

Note. *M* Model-estimated marginal mean, *CI* Confidence interval. Values are derived from the full model including time, group and the time × group interaction (random intercept for participant, REML), as shown in Figure 6. Degrees of freedom as in the coefficient tables (within-group containment method as implemented in nlme::lme; Pinheiro & Bates, 2000). Model statistics per test environment: N_participants = 36, N_observations = 72.

***Table S6 | Sequential model building and comparison for mean of temporal differences, M(Δt) (real test environment)***

| **Model** | ***df*** | ***AIC*** | ***BIC*** | ***logLik*** | **Test** | **$\chi^2$** | ***p*** | **Δ Marginal $R^2$** |
|---|---|---|---|---|---|---|---|---|
| Model 0: Intercept-only (GLS) | 2 | 749.19 | 753.74 | -372.59 | – | – | – | – |
| Model 1: Random intercept | 3 | 721.29 | 728.12 | -357.65 | 0 vs 1 | 29.90 | $p < .001$ | – |
| Model 2: + Time | 4 | 721.89 | 731.00 | -356.94 | 1 vs 2 | 1.40 | $p = .236$ | .00 |
| Model 3: + Group | 6 | 725.28 | 738.94 | -356.64 | 2 vs 3 | 0.60 | $p = .739$ | .01 |
| Model 4: + Time × Group | 8 | 722.14 | 740.36 | -353.07 | 3 vs 4 | 7.14 | $p = .028$ | .02 |

Note. *GLS* Generalized least squares, *df* Number of estimated parameters, *AIC* Akaike information criterion, *BIC* Bayesian information criterion, *logLik* Log-likelihood, $\chi^2$ Likelihood ratio test statistic against the previous model, *p* p-value, *Δ Marginal $R^2$* Change in marginal $R^2$ (variance explained by fixed effects) compared to the previous model. All models were estimated with maximum likelihood. logLik, AIC and BIC refer to the outcome in ms. An en-dash (–) indicates not applicable.

**Table S7 | Multilevel regression of mean of temporal differences, M(Δt), on time and training group in the real test environment**

| | Model 1: Random intercept | | | | | Model 4: + Time × Group | | | | |
|---|---|---|---|---|---|---|---|---|---|---|
| **Predictor** | ***B*** | ***SE*** | **95% CI** | ***t (df)*** | ***p*** | ***B*** | ***SE*** | **95% CI** | ***t (df)*** | ***p*** |
| Intercept | 15.75 | 6.77 | [2.03, 29.47] | 2.33 (36) | p = .026 | 24.90 | 12.63 | [-0.80, 50.59] | 1.97 (33) | p = .057 |
| Time | – | – | – | – | – | -22.39 | 8.08 | [-38.84, -5.95] | -2.77 (33) | p = .009 |
| Group Real | – | – | – | – | – | -12.52 | 17.86 | [-48.86, 23.82] | -0.70 (33) | p = .488 |
| Group Virtual | – | – | – | – | – | -6.07 | 17.86 | [-42.41, 30.27] | -0.34 (33) | p = .736 |
| Time × Group Real | – | – | – | – | – | 19.04 | 11.43 | [-4.21, 42.29] | 1.67 (33) | p = .105 |
| Time × Group Virtual | – | – | – | – | – | 30.43 | 11.43 | [7.18, 53.69] | 2.66 (33) | p = .012 |
| *Random effects* | | | | | | | | | | |
| Residual variance ($\sigma^2$) | 455.46 | – | – | – | – | 391.88 | – | – | – | – |
| Intercept variance ($\tau_{00}$) | 1420.14 | – | – | – | – | 1522.70 | – | – | – | – |
| ICC | .76 | – | – | – | – | .80 | – | – | – | – |
| Marginal $R^2$ | .00 | – | – | – | – | .04 | – | – | – | – |
| Conditional $R^2$ | .76 | – | – | – | – | .80 | – | – | – | – |
| Δ Marginal $R^2$ | – | – | – | – | – | .04 | – | – | – | – |
| N subjects | 36 | – | – | – | – | 36 | – | – | – | – |

Note. *B* Unstandardized regression coefficient, *SE* Standard error, *CI* Confidence interval, *t*(df) t-statistic with degrees of freedom (within-group containment method as implemented in nlme::lme; Pinheiro & Bates, 2000), *p* p-value, $\sigma^2$ Residual variance (within-subject variance), $\tau_{00}$ Random intercept variance (between-subject variance), *ICC* Intraclass correlation coefficient, *Marginal $R^2$* Variance explained by fixed effects, *Conditional $R^2$* Variance explained by fixed and random effects, *Δ Marginal $R^2$* Change in marginal $R^2$ from Model 1 to Model 4. Both models were estimated with restricted maximum likelihood (REML). Reference categories: Pre-Test for time and Solo for group. Estimates are given in ms, variances in ms². Model statistics: N_participants = 36, N_observations = 72. An en-dash (–) indicates that a value is not applicable.

***Table S8 | Sequential model building and comparison for mean of temporal differences, M(Δt) (virtual test environment)***

| **Model** | ***df*** | ***AIC*** | ***BIC*** | ***logLik*** | **Test** | **$\chi^2$** | ***p*** | **Δ Marginal $R^2$** |
|---|---|---|---|---|---|---|---|---|
| Model 0: Intercept-only (GLS) | 2 | 749.58 | 754.13 | -372.79 | – | – | – | – |
| Model 1: Random intercept | 3 | 730.12 | 736.95 | -362.06 | 0 vs 1 | 21.46 | $p < .001$ | – |
| Model 2: + Time | 4 | 732.04 | 741.15 | -362.02 | 1 vs 2 | 0.07 | $p = .787$ | .00 |
| Model 3: + Group | 6 | 735.89 | 749.55 | -361.95 | 2 vs 3 | 0.15 | $p = .927$ | .00 |
| Model 4: + Time × Group | 8 | 736.51 | 754.72 | -360.25 | 3 vs 4 | 3.39 | $p = .184$ | .01 |

Note. *GLS* Generalized least squares, *df* Number of estimated parameters, *AIC* Akaike information criterion, *BIC* Bayesian information criterion, *logLik* Log-likelihood, $\chi^2$ Likelihood ratio test statistic against the previous model, *p* p-value, *Δ Marginal $R^2$* Change in marginal $R^2$ (variance explained by fixed effects) compared to the previous model. All models were estimated with maximum likelihood. logLik, AIC and BIC refer to the outcome in ms. An en-dash (–) indicates not applicable.

**_Table S9 | Multilevel regression of mean of temporal differences, M(Δt), on time and training group in the virtual test environment_**

| | Model 1: Random intercept | | | | | Model 4: + Time × Group | | | | |
|---|---|---|---|---|---|---|---|---|---|---|
| **Predictor** | ***B*** | ***SE*** | **95% CI** | ***t (df)*** | ***p*** | ***B*** | ***SE*** | **95% CI** | ***t (df)*** | ***p*** |
| Intercept | 3.02 | 6.63 | [-10.41, 16.46] | 0.46 (36) | p = .651 | 8.44 | 12.81 | [-17.63, 34.51] | 0.66 (33) | p = .515 |
| Time | – | – | – | – | – | -11.63 | 10.01 | [-32.00, 8.74] | -1.16 (33) | p = .254 |
| Group Real | – | – | – | – | – | -15.24 | 18.12 | [-52.10, 21.62] | -0.84 (33) | p = .406 |
| Group Virtual | – | – | – | – | – | -3.36 | 18.12 | [-40.22, 33.50] | -0.19 (33) | p = .854 |
| Time × Group Real | – | – | – | – | – | 25.49 | 14.16 | [-3.31, 54.30] | 1.80 (33) | p = .081 |
| Time × Group Virtual | – | – | – | – | – | 14.09 | 14.16 | [-14.71, 42.90] | 1.00 (33) | p = .327 |
| *Random effects* | | | | | | | | | | |
| Residual variance ($\sigma^2$) | 606.88 | – | – | – | – | 601.40 | – | – | – | – |
| Intercept variance ($\tau_{00}$) | 1276.79 | – | – | – | – | 1368.26 | – | – | – | – |
| ICC | .68 | – | – | – | – | .69 | – | – | – | – |
| Marginal $R^2$ | .00 | – | – | – | – | .02 | – | – | – | – |
| Conditional $R^2$ | .68 | – | – | – | – | .70 | – | – | – | – |
| Δ Marginal $R^2$ | – | – | – | – | – | .02 | – | – | – | – |
| N subjects | 36 | – | – | – | – | 36 | – | – | – | – |

Note. *B* Unstandardized regression coefficient, *SE* Standard error, *CI* Confidence interval, *t*(df) t-statistic with degrees of freedom (within-group containment method as implemented in nlme::lme; Pinheiro & Bates, 2000), *p* p-value, $\sigma^2$ Residual variance (within-subject variance), $\tau_{00}$ Random intercept variance (between-subject variance), *ICC* Intraclass correlation coefficient, *Marginal* $R^2$ Variance explained by fixed effects, *Conditional* $R^2$ Variance explained by fixed and random effects, *Δ Marginal* $R^2$ Change in marginal $R^2$ from Model 1 to Model 4. Both models were estimated with restricted maximum likelihood (REML). Reference categories: Pre-Test for time and Solo for group. Estimates are given in ms, variances in ms². Model statistics: N_participants = 36, N_observations = 72. An en-dash (–) indicates that a value is not applicable.

***Table S10 | Sequential model building and comparison for mean of absolute acceleration differences, M(Δa) (real test environment)***

| **Model** | ***df*** | ***AIC*** | ***BIC*** | ***logLik*** | **Test** | **$\chi^2$** | ***p*** | **Δ Marginal $R^2$** |
|---|---|---|---|---|---|---|---|---|
| Model 0: Intercept-only (GLS) | 2 | -65.72 | -61.16 | 34.86 | – | – | – | – |
| Model 1: Random intercept | 3 | -120.23 | -113.40 | 63.12 | 0 vs 1 | 56.52 | p < .001 | – |
| Model 2: + Time | 4 | -118.69 | -109.58 | 63.34 | 1 vs 2 | 0.45 | p = .500 | .00 |
| Model 3: + Group | 6 | -120.91 | -107.25 | 66.45 | 2 vs 3 | 6.22 | p = .045 | .15 |
| Model 4: + Time × Group | 8 | -118.80 | -100.59 | 67.40 | 3 vs 4 | 1.90 | p = .387 | .00 |

Note. *GLS* Generalized least squares, *df* Number of estimated parameters, *AIC* Akaike information criterion, *BIC* Bayesian information criterion, *logLik* Log-likelihood, $\chi^2$ Likelihood ratio test statistic against the previous model, *p* p-value, *Δ Marginal $R^2$* Change in marginal $R^2$ (variance explained by fixed effects) compared to the previous model. All models were estimated with maximum likelihood. An en-dash (–) indicates not applicable.

### *Table S11 | Multilevel regression of mean of absolute acceleration differences, M(Δa), on time and training group in the real test environment*

| | *Model 1: Random intercept* | | | | | *Model 4: + Time × Group* | | | | |
|---|---|---|---|---|---|---|---|---|---|---|
| **Predictor** | ***B*** | ***SE*** | **95% CI** | ***t (df)*** | ***p*** | ***B*** | ***SE*** | **95% CI** | ***t (df)*** | ***p*** |
| Intercept | 0.06 | 0.02 | [0.01, 0.11] | 2.28 (36) | p = .029 | 0.13 | 0.04 | [0.04, 0.21] | 3.05 (33) | p = .005 |
| Time | – | – | – | – | – | 0.01 | 0.02 | [-0.03, 0.05] | 0.63 (33) | p = .530 |
| Group Real | – | – | – | – | – | -0.07 | 0.06 | [-0.19, 0.05] | -1.22 (33) | p = .231 |
| Group Virtual | – | – | – | – | – | -0.13 | 0.06 | [-0.25, -0.01] | -2.18 (33) | p = .037 |
| Time × Group Real | – | – | – | – | – | -0.04 | 0.03 | [-0.10, 0.02] | -1.32 (33) | p = .196 |
| Time × Group Virtual | – | – | – | – | – | -0.02 | 0.03 | [-0.08, 0.03] | -0.84 (33) | p = .407 |
| *Random effects* | | | | | | | | | | |
| Residual variance ($\sigma^2$) | 0.00 | – | – | – | – | 0.00 | – | – | – | – |
| Intercept variance ($\tau_{00}$) | 0.02 | – | – | – | – | 0.02 | – | – | – | – |
| ICC | .89 | – | – | – | – | .88 | – | – | – | – |
| Marginal $R^2$ | .00 | – | – | – | – | .14 | – | – | – | – |
| Conditional $R^2$ | .89 | – | – | – | – | .90 | – | – | – | – |
| Δ Marginal $R^2$ | – | – | – | – | – | .14 | – | – | – | – |
| N subjects | 36 | – | – | – | – | 36 | – | – | – | – |

Note. *B* Unstandardized regression coefficient, *SE* Standard error, *CI* Confidence interval, *t*(df) t-statistic with degrees of freedom (within-group containment method as implemented in nlme::lme; Pinheiro & Bates, 2000), *p* p-value, $\sigma^2$ Residual variance (within-subject variance), $\tau_{00}$ Random intercept variance (between-subject variance), *ICC* Intraclass correlation coefficient, *Marginal* $R^2$ Variance explained by fixed effects, *Conditional* $R^2$ Variance explained by fixed and random effects, *Δ Marginal* $R^2$ Change in marginal $R^2$ from Model 1 to Model 4. Both models were estimated with restricted maximum likelihood (REML). Reference categories: Pre-Test for time and Solo for group. Model statistics: N_participants = 36, N_observations = 72. An en-dash (–) indicates that a value is not applicable.

***Table S12 | Sequential model building and comparison for mean of absolute acceleration differences, M(Δa) (virtual test environment)***

| **Model** | ***df*** | ***AIC*** | ***BIC*** | ***logLik*** | **Test** | **$\chi^2$** | ***p*** | **Δ Marginal $R^2$** |
|---|---|---|---|---|---|---|---|---|
| Model 0: Intercept-only (GLS) | 2 | -78.28 | -73.72 | 41.14 | – | – | – | – |
| Model 1: Random intercept | 3 | -126.95 | -120.12 | 66.48 | 0 vs 1 | 50.68 | $p < .001$ | – |
| Model 2: + Time | 4 | -125.51 | -116.40 | 66.75 | 1 vs 2 | 0.55 | $p = .457$ | .00 |
| Model 3: + Group | 6 | -124.70 | -111.04 | 68.35 | 2 vs 3 | 3.19 | $p = .203$ | .08 |
| Model 4: + Time × Group | 8 | -124.03 | -105.82 | 70.02 | 3 vs 4 | 3.34 | $p = .188$ | .01 |

Note. *GLS* Generalized least squares, *df* Number of estimated parameters, *AIC* Akaike information criterion, *BIC* Bayesian information criterion, *logLik* Log-likelihood, $\chi^2$ Likelihood ratio test statistic against the previous model, *p* p-value, *Δ Marginal $R^2$* Change in marginal $R^2$ (variance explained by fixed effects) compared to the previous model. All models were estimated with maximum likelihood. An en-dash (–) indicates not applicable.

***Table S13 | Multilevel regression of mean of absolute acceleration differences, M(Δa), on time and training group in the virtual test environment***

| | Model 1: Random intercept | | | | | Model 4: + Time × Group | | | | |
|---|---|---|---|---|---|---|---|---|---|---|
| **Predictor** | ***B*** | ***SE*** | **95% CI** | ***t (df)*** | ***p*** | ***B*** | ***SE*** | **95% CI** | ***t (df)*** | ***p*** |
| Intercept | 0.06 | 0.02 | [0.02, 0.11] | 2.80 (36) | p = .008 | 0.10 | 0.04 | [0.02, 0.18] | 2.64 (33) | p = .013 |
| Time | – | – | – | – | – | 0.02 | 0.02 | [-0.02, 0.06] | 0.97 (33) | p = .341 |
| Group Real | – | – | – | – | – | -0.04 | 0.06 | [-0.16, 0.07] | -0.79 (33) | p = .436 |
| Group Virtual | – | – | – | – | – | -0.07 | 0.06 | [-0.18, 0.05] | -1.22 (33) | p = .231 |
| Time × Group Real | – | – | – | – | – | -0.03 | 0.03 | [-0.09, 0.02] | -1.22 (33) | p = .231 |
| Time × Group Virtual | – | – | – | – | – | -0.05 | 0.03 | [-0.11, 0.01] | -1.75 (33) | p = .090 |
| *Random effects* | | | | | | | | | | |
| Residual variance ($\sigma^2$) | 0.00 | – | – | – | – | 0.00 | – | – | – | – |
| Intercept variance ($\tau_{00}$) | 0.02 | – | – | – | – | 0.02 | – | – | – | – |
| ICC | .87 | – | – | – | – | .87 | – | – | – | – |
| Marginal $R^2$ | .00 | – | – | – | – | .08 | – | – | – | – |
| Conditional $R^2$ | .87 | – | – | – | – | .88 | – | – | – | – |
| Δ Marginal $R^2$ | – | – | – | – | – | .08 | – | – | – | – |
| N subjects | 36 | – | – | – | – | 36 | – | – | – | – |

Note. *B* Unstandardized regression coefficient, *SE* Standard error, *CI* Confidence interval, *t*(df) t-statistic with degrees of freedom (within-group containment method as implemented in nlme::lme; Pinheiro & Bates, 2000), *p* p-value, $\sigma^2$ Residual variance (within-subject variance), $\tau_{00}$ Random intercept variance (between-subject variance), *ICC* Intraclass correlation coefficient, *Marginal $R^2$* Variance explained by fixed effects, *Conditional $R^2$* Variance explained by fixed and random effects, *Δ Marginal $R^2$* Change in marginal $R^2$ from Model 1 to Model 4. Both models were estimated with restricted maximum likelihood (REML). Reference categories: Pre-Test for time and Solo for group. Model statistics: N_participants = 36, N_observations = 72. An en-dash (–) indicates that a value is not applicable.

***Table S14 | Sequential model building and comparison for dispersion of absolute acceleration differences, SD(Δa) (real test environment)***

| Model | *df* | *AIC* | *BIC* | *logLik* | Test | $\chi^2$ | *p* | Δ Marginal $R^2$ |
|---|---|---|---|---|---|---|---|---|
| Model 0: Intercept-only (GLS) | 2 | -285.13 | -280.58 | 144.56 | – | – | – | – |
| Model 1: Random intercept | 3 | -284.06 | -277.23 | 145.03 | 0 vs 1 | 0.93 | p = .334 | – |
| Model 2: + Time | 4 | -288.96 | -279.85 | 148.48 | 1 vs 2 | 6.90 | p = .009 | .07 |
| Model 3: + Group | 6 | -300.37 | -286.71 | 156.19 | 2 vs 3 | 15.41 | p < .001 | .20 |
| Model 4: + Time × Group | 8 | -297.54 | -279.33 | 156.77 | 3 vs 4 | 1.17 | p = .557 | .01 |

Note. *GLS* Generalized least squares, *df* Number of estimated parameters, *AIC* Akaike information criterion, *BIC* Bayesian information criterion, *logLik* Log-likelihood, $\chi^2$ Likelihood ratio test statistic against the previous model, *p* p-value, *Δ Marginal* $R^2$ Change in marginal $R^2$ (variance explained by fixed effects) compared to the previous model. All models were estimated with maximum likelihood. An en-dash (–) indicates not applicable.

***Table S15 | Multilevel regression of dispersion of absolute acceleration differences, SD(Δa), on time and training group in the real test environment***

| | Model 3: + Group | | | | | Model 4: + Time × Group | | | | |
|---|---|---|---|---|---|---|---|---|---|---|
| **Predictor** | ***B*** | ***SE*** | **95% CI** | ***t (df)*** | ***p*** | ***B*** | ***SE*** | **95% CI** | ***t (df)*** | ***p*** |
| Intercept | 0.14 | 0.01 | [0.13, 0.16] | 20.87 (35) | p < .001 | 0.14 | 0.01 | [0.12, 0.15] | 16.66 (33) | p < .001 |
| Time | -0.02 | 0.01 | [-0.03, -0.00] | -2.72 (35) | p = .010 | -0.01 | 0.01 | [-0.03, 0.02] | -0.71 (33) | p = .483 |
| Group Real | 0.03 | 0.01 | [0.01, 0.04] | 3.11 (33) | p = .004 | 0.03 | 0.01 | [0.01, 0.06] | 2.94 (33) | p = .006 |
| Group Virtual | 0.03 | 0.01 | [0.02, 0.05] | 4.00 (33) | p < .001 | 0.04 | 0.01 | [0.02, 0.06] | 3.46 (33) | p = .002 |
| Time × Group Real | – | – | – | – | – | -0.02 | 0.02 | [-0.05, 0.02] | -0.98 (33) | p = .332 |
| Time × Group Virtual | – | – | – | – | – | -0.01 | 0.02 | [-0.05, 0.02] | -0.80 (33) | p = .432 |
| *Random effects* | | | | | | | | | | |
| Residual variance ($\sigma^2$) | 0.00 | – | – | – | – | 0.00 | – | – | – | – |
| Intercept variance ($\tau_{00}$) | 0.00 | – | – | – | – | 0.00 | – | – | – | – |
| ICC | .07 | – | – | – | – | .06 | – | – | – | – |
| Marginal $R^2$ | .27 | – | – | – | – | .27 | – | – | – | – |
| Conditional $R^2$ | .32 | – | – | – | – | .31 | – | – | – | – |
| Δ Marginal $R^2$ | – | – | – | – | – | .01 | – | – | – | – |
| N subjects | 36 | – | – | – | – | 36 | – | – | – | – |

Note. *B* Unstandardized regression coefficient, *SE* Standard error, *CI* Confidence interval, *t*(df) t-statistic with degrees of freedom (within-group containment method as implemented in nlme::lme; Pinheiro & Bates, 2000), *p* p-value, $\sigma^2$ Residual variance (within-subject variance), $\tau_{00}$ Random intercept variance (between-subject variance), *ICC* Intraclass correlation coefficient, *Marginal $R^2$* Variance explained by fixed effects, *Conditional $R^2$* Variance explained by fixed and random effects, *Δ Marginal $R^2$* Change in marginal $R^2$ from Model 3 to Model 4. Both models were estimated with restricted maximum likelihood (REML). Reference categories: Pre-Test for time and Solo for group. Model statistics: N_participants = 36, N_observations = 72. An en-dash (–) indicates that a value is not applicable.

***Table S16 | Sequential model building and comparison for dispersion of absolute acceleration differences, SD(Δa) (virtual test environment)***

| **Model** | ***df*** | ***AIC*** | ***BIC*** | ***logLik*** | **Test** | $\chi^2$ | ***p*** | **Δ Marginal R²** |
|---|---|---|---|---|---|---|---|---|
| Model 0: Intercept-only (GLS) | 2 | -290.99 | -286.44 | 147.49 | – | – | – | – |
| Model 1: Random intercept | 3 | -295.47 | -288.64 | 150.73 | 0 vs 1 | 6.48 | p = .011 | – |
| Model 2: + Time | 4 | -312.87 | -303.77 | 160.44 | 1 vs 2 | 19.41 | p < .001 | .13 |
| Model 3: + Group | 6 | -317.98 | -304.32 | 164.99 | 2 vs 3 | 9.11 | p = .011 | .16 |
| Model 4: + Time × Group | 8 | -315.91 | -297.70 | 165.96 | 3 vs 4 | 1.93 | p = .381 | .01 |

Note. *GLS* Generalized least squares, *df* Number of estimated parameters, *AIC* Akaike information criterion, *BIC* Bayesian information criterion, *logLik* Log-likelihood, $\chi^2$ Likelihood ratio test statistic against the previous model, *p* p-value, *Δ Marginal R²* Change in marginal $R^2$ (variance explained by fixed effects) compared to the previous model. All models were estimated with maximum likelihood. An en-dash (–) indicates not applicable.

***Table S17 | Multilevel regression of dispersion of absolute acceleration differences, SD(Δa), on time and training group in the virtual test environment***

| | Model 3: + Group | | | | | Model 4: + Time × Group | | | | |
|---|---|---|---|---|---|---|---|---|---|---|
| **Predictor** | ***B*** | ***SE*** | **95% CI** | ***t (df)*** | ***p*** | ***B*** | ***SE*** | **95% CI** | ***t (df)*** | ***p*** |
| Intercept | 0.16 | 0.01 | [0.14, 0.17] | 21.28 (35) | p < .001 | 0.15 | 0.01 | [0.14, 0.17] | 19.51 (33) | p < .001 |
| Time | -0.02 | 0.00 | [-0.03, -0.01] | -5.00 (35) | p < .001 | -0.02 | 0.01 | [-0.04, -0.01] | -2.77 (33) | p = .009 |
| Group Real | 0.03 | 0.01 | [0.01, 0.05] | 2.64 (33) | p = .012 | 0.02 | 0.01 | [0.00, 0.05] | 2.05 (33) | p = .048 |
| Group Virtual | 0.03 | 0.01 | [0.01, 0.05] | 2.70 (33) | p = .011 | 0.03 | 0.01 | [0.01, 0.05] | 2.74 (33) | p = .010 |
| Time × Group Real | – | – | – | – | – | 0.01 | 0.01 | [-0.02, 0.03] | 0.55 (33) | p = .586 |
| Time × Group Virtual | – | – | – | – | – | -0.01 | 0.01 | [-0.03, 0.01] | -0.79 (33) | p = .435 |
| *Random effects* | | | | | | | | | | |
| Residual variance ($\sigma^2$) | 0.00 | – | – | – | – | 0.00 | – | – | – | – |
| Intercept variance ($\tau_{00}$) | 0.00 | – | – | – | – | 0.00 | – | – | – | – |
| ICC | .54 | – | – | – | – | .54 | – | – | – | – |
| Marginal $R^2$ | .27 | – | – | – | – | .28 | – | – | – | – |
| Conditional $R^2$ | .66 | – | – | – | – | .66 | – | – | – | – |
| Δ Marginal $R^2$ | – | – | – | – | – | .01 | – | – | – | – |
| N subjects | 36 | – | – | – | – | 36 | – | – | – | – |

Note. *B* Unstandardized regression coefficient, *SE* Standard error, *CI* Confidence interval, *t*(df) t-statistic with degrees of freedom (within-group containment method as implemented in nlme::lme; Pinheiro & Bates, 2000), *p* p-value, $\sigma^2$ Residual variance (within-subject variance), $\tau_{00}$ Random intercept variance (between-subject variance), *ICC* Intraclass correlation coefficient, *Marginal $R^2$* Variance explained by fixed effects, *Conditional $R^2$* Variance explained by fixed and random effects, *Δ Marginal $R^2$* Change in marginal $R^2$ from Model 3 to Model 4. Both models were estimated with restricted maximum likelihood (REML). Reference categories: Pre-Test for time and Solo for group. Model statistics: N_participants = 36, N_observations = 72. An en-dash (–) indicates that a value is not applicable.

### *Table S18 | Sequential model building and comparison for the combined data of both test environments*

| Model | *df* | *AIC* | *BIC* | *logLik* | Test | $\chi^2$ | *p* | Δ Marginal $R^2$ |
|---|---|---|---|---|---|---|---|---|
| *Dispersion of temporal differences, SD(Δt)* | | | | | | | | |
| Model 0: Intercept-only (GLS) | 2 | 1407.10 | 1413.04 | -701.55 | – | – | – | – |
| Model 1: Random intercept | 3 | 1386.19 | 1395.10 | -690.09 | 0 vs 1 | 22.91 | p < .001 | – |
| Model 2: + Time | 4 | 1360.65 | 1372.53 | -676.33 | 1 vs 2 | 27.54 | p < .001 | .11 |
| Model 3: + Group | 6 | 1359.95 | 1377.76 | -673.97 | 2 vs 3 | 4.71 | p = .095 | .06 |
| Model 4: + Time × Group | 8 | 1363.49 | 1387.25 | -673.74 | 3 vs 4 | 0.46 | p = .795 | .00 |
| Model 5: + Environment | 9 | 1361.96 | 1388.69 | -671.98 | 4 vs 5 | 3.53 | p = .060 | .01 |
| Model 6: + Environment × Time | 10 | 1361.01 | 1390.71 | -670.51 | 5 vs 6 | 2.95 | p = .086 | .01 |
| Model 7: + Environment × Group | 12 | 1357.05 | 1392.69 | -666.53 | 6 vs 7 | 7.96 | p = .019 | .02 |
| Model 8: + Time × Group × Environment | 14 | 1358.42 | 1400.00 | -665.21 | 7 vs 8 | 2.64 | p = .268 | .01 |
| R: Retained model (random intercept + time) | 4 | 1360.65 | 1372.53 | -676.33 | – | – | – | – |
| R + E: Retained model + Environment | 5 | 1359.14 | 1373.99 | -674.57 | R vs R + E | 3.51 | p = .061 | .01 |
| *Mean of temporal differences, M(Δt)* | | | | | | | | |
| Model 0: Intercept-only (GLS) | 2 | 1497.91 | 1503.85 | -746.96 | – | – | – | – |
| Model 1: Random intercept | 3 | 1417.85 | 1426.76 | -705.93 | 0 vs 1 | 82.06 | p < .001 | – |
| Model 2: + Time | 4 | 1419.57 | 1431.45 | -705.79 | 1 vs 2 | 0.28 | p = .597 | .00 |
| Model 3: + Group | 6 | 1423.20 | 1441.01 | -705.60 | 2 vs 3 | 0.38 | p = .829 | .01 |
| Model 4: + Time × Group | 8 | 1420.43 | 1444.19 | -702.21 | 3 vs 4 | 6.77 | p = .034 | .01 |
| Model 5: + Environment | 9 | 1411.64 | 1438.37 | -696.82 | 4 vs 5 | 10.78 | p = .001 | .02 |
| Model 6: + Environment × Time | 10 | 1412.66 | 1442.36 | -696.33 | 5 vs 6 | 0.98 | p = .322 | .00 |
| Model 7: + Environment × Group | 12 | 1416.15 | 1451.78 | -696.07 | 6 vs 7 | 0.52 | p = .773 | .00 |
| Model 8: + Time × Group × Environment | 14 | 1418.50 | 1460.08 | -695.25 | 7 vs 8 | 1.65 | p = .439 | .00 |
| R: Retained model (random intercept only) | 3 | 1417.85 | 1426.76 | -705.93 | – | – | – | – |
| R + E: Retained model + Environment | 4 | 1409.78 | 1421.66 | -700.89 | R vs R + E | 10.07 | p = .002 | .02 |

| Model | *df* | *AIC* | *BIC* | *logLik* | Test | $\chi^2$ | *p* | Δ Marginal $R^2$ |
|---|---|---|---|---|---|---|---|---|
| *Mean of absolute acceleration differences, M(Δa)* | | | | | | | | |
| Model 0: Intercept-only (GLS) | 2 | -147.37 | -141.43 | 75.69 | – | – | – | – |
| Model 1: Random intercept | 3 | -338.69 | -329.78 | 172.34 | 0 vs 1 | 193.32 | $p < .001$ | – |
| Model 2: + Time | 4 | -337.79 | -325.92 | 172.90 | 1 vs 2 | 1.11 | $p = .293$ | .00 |
| Model 3: + Group | 6 | -338.52 | -320.70 | 175.26 | 2 vs 3 | 4.73 | $p = .094$ | .11 |
| Model 4: + Time × Group | 8 | -339.51 | -315.75 | 177.75 | 3 vs 4 | 4.99 | $p = .083$ | .00 |
| Model 5: + Environment | 9 | -338.26 | -311.53 | 178.13 | 4 vs 5 | 0.75 | $p = .385$ | .00 |
| Model 6: + Environment × Time | 10 | -336.26 | -306.57 | 178.13 | 5 vs 6 | 0.00 | $p = .958$ | .00 |
| Model 7: + Environment × Group | 12 | -338.91 | -303.27 | 181.45 | 6 vs 7 | 6.64 | $p = .036$ | .00 |
| Model 8: + Time × Group × Environment | 14 | -335.66 | -294.08 | 181.83 | 7 vs 8 | 0.75 | $p = .686$ | .00 |
| R: Retained model (random intercept only) | 3 | -338.69 | -329.78 | 172.34 | – | – | – | – |
| R + E: Retained model + Environment | 4 | -337.40 | -325.52 | 172.70 | R vs R + E | 0.71 | $p = .399$ | .00 |
| *Dispersion of absolute acceleration differences, SD(Δa)* | | | | | | | | |
| Model 0: Intercept-only (GLS) | 2 | -578.01 | -572.07 | 291.01 | – | – | – | – |
| Model 1: Random intercept | 3 | -597.79 | -588.88 | 301.89 | 0 vs 1 | 21.77 | $p < .001$ | – |
| Model 2: + Time | 4 | -619.59 | -607.71 | 313.79 | 1 vs 2 | 23.80 | $p < .001$ | .10 |
| Model 3: + Group | 6 | -630.38 | -612.56 | 321.19 | 2 vs 3 | 14.80 | $p < .001$ | .18 |
| Model 4: + Time × Group | 8 | -627.68 | -603.92 | 321.84 | 3 vs 4 | 1.30 | $p = .522$ | .00 |
| Model 5: + Environment | 9 | -629.64 | -602.91 | 323.82 | 4 vs 5 | 3.95 | $p = .047$ | .01 |
| Model 6: + Environment × Time | 10 | -627.98 | -598.28 | 323.99 | 5 vs 6 | 0.34 | $p = .559$ | .00 |
| Model 7: + Environment × Group | 12 | -624.83 | -589.20 | 324.42 | 6 vs 7 | 0.86 | $p = .652$ | .00 |
| Model 8: + Time × Group × Environment | 14 | -622.44 | -580.87 | 325.22 | 7 vs 8 | 1.61 | $p = .447$ | .01 |
| R: Retained model (random intercept + time + group) | 6 | -630.38 | -612.56 | 321.19 | – | – | – | – |
| R + E: Retained model + Environment | 7 | -632.29 | -611.50 | 323.14 | R vs R + E | 3.91 | $p = .048$ | .01 |

| Model | *df* | *AIC* | *BIC* | *logLik* | Test | *χ²* | *p* | Δ Marginal R² |
|---|---|---|---|---|---|---|---|---|

Note. Data of both test environments combined (N_participants = 36, N_observations = 144). Environment = test environment (real vs. virtual). GLS Generalized least squares, df Number of estimated parameters, AIC Akaike information criterion, BIC Bayesian information criterion, logLik Log-likelihood, $\chi^2$ Likelihood ratio test statistic against the previous model, p p-value, Δ Marginal $R^2$ Change in marginal $R^2$ (variance explained by fixed effects) compared to the previous model. All models were estimated with maximum likelihood. For SD(Δt) and M(Δt), logLik, AIC and BIC refer to the outcome in ms. Rows R and R + E: retained model of the separate analyses, compared without and with environment as additional fixed effect. An en-dash (–) indicates not applicable.
