## Additional file 1 for "Joint Walking Training with Virtual and Real Partners: Testing the Transfer from Virtual Reality to the Real World"

**S1: English version of the VEQ and additional questions translated**

Please read each statement and check the relevant response to indicate how strongly you agree or disagree with each statement (1 through 7). There are no right or wrong answers. Answer spontaneously and intuitively.
The first questions refer to your experiences with your own body in the virtual world.

| **Acceptance/ Body Ownership** | Strongly disagree 1 | Disagree 2 | Somewhat disagree 3 | Neither agree nor disagree 4 | Somewhat agree 5 | Agree 6 | Strongly agree 7 |
|---|---|---|---|---|---|---|---|
| It felt like the virtual body was my body. | 0 | 0 | 0 | 0 | 0 | 0 | 0 |
| It felt like the virtual body parts were my body parts. | 0 | 0 | 0 | 0 | 0 | 0 | 0 |
| The virtual body felt like a human body. | 0 | 0 | 0 | 0 | 0 | 0 | 0 |
| It felt like the virtual body belonged to me. | 0 | 0 | 0 | 0 | 0 | 0 | 0 |

| **Control/ Agency** | Strongly disagree 1 | Disagree 2 | Somewhat disagree 3 | Neither agree nor disagree 4 | Somewhat agree 5 | Agree 6 | Strongly agree 7 |
|---|---|---|---|---|---|---|---|
| The movements of the virtual body felt like they were my movements. | 0 | 0 | 0 | 0 | 0 | 0 | 0 |
| I felt like I was controlling the movements of the virtual body. | 0 | 0 | 0 | 0 | 0 | 0 | 0 |
| I felt like I was causing the movements of the virtual body. | 0 | 0 | 0 | 0 | 0 | 0 | 0 |
| The movements of the virtual body were in sync with my own movements. | 0 | 0 | 0 | 0 | 0 | 0 | 0 |

| Change | Strongly disagree<br>1 | Disagree<br>2 | Somewhat disagree<br>3 | Neither agree nor disagree<br>4 | Somewhat agree<br>5 | Agree<br>6 | Strongly agree<br>7 |
|---|---|---|---|---|---|---|---|
| I felt like the form or appearance of my own body had changed. | O | O | O | O | O | O | O |
| I felt like the weight of my own body had changed. | O | O | O | O | O | O | O |
| I felt like the size (height) of my own body had changed. | O | O | O | O | O | O | O |
| I felt like the width of my own body had changed. | O | O | O | O | O | O | O |

The next questions refer to your perception of the leader. Please again check to what extent you agree with them.

| **Perception of the leader** | Strongly disagree<br>1 | Disagree<br>2 | Somewhat disagree<br>3 | Neither agree nor disagree<br>4 | Somewhat agree<br>5 | Agree<br>6 | Strongly agree<br>7 |
|---|---|---|---|---|---|---|---|
| It felt like the virtual leader was a real person. | 0 | 0 | 0 | 0 | 0 | 0 | 0 |
| It felt like the virtual body parts of the leader were those of a real human. | 0 | 0 | 0 | 0 | 0 | 0 | 0 |
| The virtual body of the leader felt like a human body. | 0 | 0 | 0 | 0 | 0 | 0 | 0 |
| The virtual leader felt real. | 0 | 0 | 0 | 0 | 0 | 0 | 0 |
| The movements of the virtual leader were realistic. | 0 | 0 | 0 | 0 | 0 | 0 | 0 |
| For me, it was more difficult to follow the leader in the virtual world than in the real world. | 0 | 0 | 0 | 0 | 0 | 0 | 0 |
| I felt like the leader in front of me had the same body in the virtual world as in the real world. | 0 | 0 | 0 | 0 | 0 | 0 | 0 |
